# A Normative Bayesian Model of Classification for Agents with Bounded Memory

**DOI:** 10.1101/787424

**Authors:** Heeseung Lee, Hyang-Jung Lee, Kyoung Whan Choe, Sang-Hun Lee

## Abstract

Classification, one of the key ingredients for human cognition, entails establishing a criterion that splits a given feature space into mutually exclusive subspaces. In classification tasks performed in daily life, however, a criterion is often not provided explicitly but instead needs to be guessed from past samples of a feature space. For example, we judge today’s temperature to be “cold” or “warm” by implicitly comparing it against a “typical” seasonal temperature. In such situations, establishing an optimal criterion is challenging for cognitive agents with bounded memory because it requires retrieving an entire set of past episodes with precision. As a computational account for how humans carry out this challenging operation, we developed a normative Bayesian model of classification (NBMC), in which Bayesian agents, whose working-memory precision decays as episodes elapse, continuously update their criterion as they perform a binary perceptual classification task on sequentially presented stimuli. We drew a set of specific implications regarding key properties of classification from the NBMC, and demonstrated the correspondence between the NBMC and human observers in classification behavior for each of those implications. Furthermore, in the functional magnetic resonance imaging responses acquired concurrently with behavioral data, we identified an ensemble of brain activities that coherently represent the latent variables, including the inferred criterion, of the NBMC. Given these results, we believe that the NBMC is worth being considered as a useful computational model that guides behavioral and neural studies on perceptual classification, especially for agents with bounded memory representation of past sensory events.

**Significant Statement:** Although classification—assigning events into mutually exclusive classes—requires a criterion, people often have to perform various classification tasks without explicit criteria. In such situations, forming a criterion based on past experience is quite challenging because people’s memory of past events deteriorates quickly over time. Here, we provided a computational model for how a memory-bounded yet normative agent infers the criterion from past episodes to maximally perform a binary perceptual classification task. This model successfully captured several key properties of human classification behavior, and the neural signals representing its latent variables were identified in the classifying human brains. By offering a rational account for memory-bonded agents’ classification, our model can guide future behavioral and neural studies on perceptual classification.

## Introduction

Classification, an act of organizing entities into mutually exclusive classes according to a criterion, is an essential ingredient of human cognition. Classification is necessary to the emergence of scientific constructs such as taxonomies in biology (Hempel, 1965; Jacob, 2004) and entailed in generating basic linguistic propositions such as predication (Rips and Turnbull, 1980), being frequently exercised when describing daily events (e.g., ‘the train arrived earlier/later than the scheduled time’).

A criterion is necessary for classification but often not explicitly provided. Previous modelling efforts offered several algorithms for how a criterion is guessed in such situations (Lapid et al., 2008; Raviv et al., 2012; Treisman and Williams, 1984) and described human classification behavior (Lages and Treisman, 1998). However, they did not specify any process of minimizing classification errors, leaving it unclear how their algorithms can be adopted from the optimality perspective. Furthermore, they did not specify how uncertainty—which prevails in the world and is critical for normative information processing (Gorgoraptis et al., 2011; Kersten et al., 2004; Körding and Wolpert, 2004; Zokaei et al., 2015)—is incorporated into their algorithms. Here, to overcome these limitations, we propose a normative Bayesian model of classification (NBMC) with an explicit cost function and probabilistic representation of uncertainty (Griffiths et al., 2012; Ma, 2012).

For model transparency and validation, we developed the NBMC based on a simple perceptual classification task^1^. This task requires classifying sequentially presented stimuli into ‘large’ or ‘small’ sizes with respect to their median size, which is not explicitly presented to observers (Fig. 1). Given this statistical structure, the best criterion computation is to infer the median size of the stimuli that have been experienced so far from the noisy sensory evidences instigated by those stimuli. Critically, we assumed that these sensory evidences become increasingly uncertain as trials elapse, following recent reports on the temporal decay of human working memory precision (Gorgoraptis et al., 2011; Zokaei et al., 2015). Accordingly, the NBMC modeled humans as Bayesian observers who infer the criterion from past sensory evidences by incorporating the temporal increase of memory uncertainty into their generative models. Then, by inferring the size of a current stimulus (s) and comparing that against the inferred criterion (c), the Bayesian observers make a decision.

**Figure 1.**
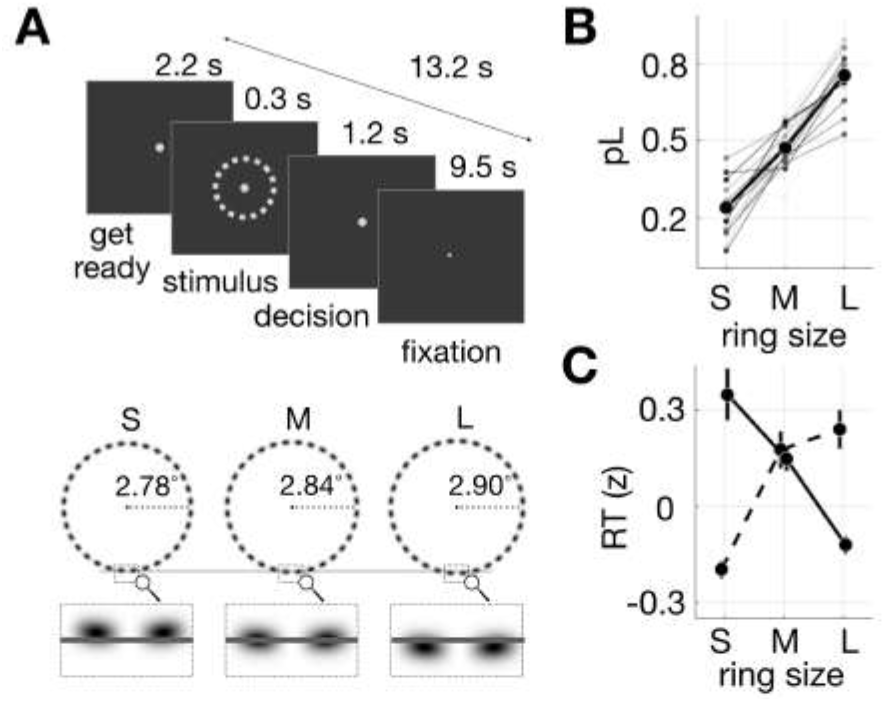
Task and behavioral measures. ***A,*** Ring-size classification task. On each trial, observers briefly viewed a ring stimulus and classified it as ‘small’ or ‘large’ within 1.2 s after stimulus onset while fixating on a spot on the display (top). Three ring stimuli that were slightly different in size (bottom) were presented in a random order. The luminance polarity of the ring images at the bottom is reversed here for illustrative purposes. ***B,*** Proportion of ‘large’ decisions (pL) as a function of ring size. Thin and bold markers, pLs for the individual observers and across-observer average, respectively. ***C,*** Z-scored RT plotted against ring size. Dashed and solid lines represent data from trials on which ‘small’ and ‘large’ decisions were made, respectively. *B*, *C*, Circles and error bars indicate the mean and standard error of the mean across observers, respectively. The stimuli, procedure, and data from the main data set (see Methods and Table 1) are shown here.

The NBMC has the following useful (Kording et al., 2020) implications regarding the key properties of perceptual classification. Firstly, the NBMC implicates specific history effects: (i) only past stimuli, but not past decisions, bias current decisions because c varies only as a function of past sensory evidences; (ii) this stimulus history effect will diminish as trials elapse because the Bayes-optimal integration weights for sensory evidences decrease as the degree of uncertainty increases (Ernst and Banks, 2002; Körding and Wolpert, 2004). Secondly, because classification outcomes are determined by the interplay between s and c, (i) the trial-to-trial variability of decision and its uncertainty can be dissected into those from two distinct sources, s and c, and (ii) classification outcomes are not governed by absolute quantities of stimuli but by relative quantities between s and c. Lastly, because the decision variable (v) for classification is determined by the difference between s and c, these three latent variables of the NBMC constitute the causal structure of ‘common effect’ (s → v ← c). This implicates that (i) there exist the respective signals of s, c, and v in the human brain engaged in classification and that (ii) the correlations among those brain signals are consistent with the causal structure prescribed by the NBMC. We verified these implications by examining the behavioral and functional-magnetic-resonance-imaging (fMRI) brain responses from human observers performing a perceptual classification task.

**Table 1.** Specifications of the data sets for their origins and experimental procedures.

| Data set # | DS#1<br>(Main data) | DS#2 | DS#3 | DS#4 |
| --- | --- | --- | --- | --- |
| Presented previously | No | Yes | No | Yes |
| Source | - | Choe et al., 2014; paper; Choe et al., 2016; paper | Choe et al., 2014; paper; Choe et al., 2016; paper | Lee et al., 2016; Conference poster |
| Feedback interval | Run-to-run | Run-to-run | Trial-to-trial | Trial-to-trial |
| Feedback manipulation | Deterministic | Deterministic | Deterministic | Stochastic |
| Inter-trial interval (s) | 13.2 | 13.2 | 2 | 2.5 |
| The number of stimuli | 3 | 3 | 16 | 5 |
| Participants number | 18 | 23 | 23 | 30 |
| With fMRI imaging | Yes | No | No | No |
| Training sessions | Yes | No | No | No |

## Materials and Methods

A total of four data sets was acquired (Table 1). The main data set (DS#1) was acquired from 18 participants (9 females, aged 20–30 years), who performed the task inside an MRI scanner. The current sample size is comparable to that used in previous fMRI studies with methods similar to the current study (Kahnt et al., 2011a; Kahnt et al., 2011b). The remaining three data sets (DS#2~4) were borrowed from previously published (Choe et al., 2014, 2016) or in-preparation (Choe et al., 2014, 2016; Lee et al., 2016) papers in our lab. DS#1 was never published nor used in any previous work. The Research Ethics Committee of Seoul National University approved the experimental procedures. All participants gave informed consent and were naïve to the purpose of the experiments.

### Behavioral data acquisition for DS#1

Figure 1*A* illustrates the experimental procedure and stimuli for DS#1. While fixating at the screen center, observers were instructed to view a brief (0.3s) ring-shape stimulus and classify its size within 1.5s after stimulus onset into either ‘small’ or ‘large’ by pressing the left-hand and right-hand keys, respectively. The identity (left/right) and timing of each key press were recorded.

Task conditions for the training (pre-scanning) sessions. Before participating in the main fMRI experiment, observers had practiced on the task intensively over several (3 to 6) training sessions (~1,000 trials for each session) outside the scanner until they reached an asymptotic level in accuracy. On an initial block of trials (~400) of each training session, we presented ring stimuli of 24 different, fine-grained radii (7.65° ~ 10.35°) in the order prescribed by an adaptive staircase (1-up-2-down) method and provided oberservers with trial-to-trial feedback based on a fixed, objective classification criterion (radius of 9°) to help them understand the goal of the task (i.e., to maximize the proportion of correct classification) and to determine the three threshold-level ring sizes tailored for a given observer. Then, on the following block of trials (~600), we asked observers to perform the task on this tailored triplet of ring stimuli while providing them with run-to-run feedback to help them get used to the task conditions under which they will later perform the task inside the MR scanner. During the training sessions, consecutive trials were apart from one another by 2.7s. Note that we opted to train observers with the stimuli that were much larger from those for the main experiment (mean radius of 2.84°) to avoid any unwanted adaptation or learning effects at low sensory levels and thus to focus training on the task structure of perceptual classification.

Task conditions for the main (scanning) session. The task conditions for the main session were identical to those for the training sessions, except for the following. Consecutive trials were apart from one another by 13.2s. The long inter-stimulus interval in conjunction with the brief stimulus presentation was implemented to minimize possible sensory adaptation to stimuli, and thus to prevent sensory adaptation in previous trials from interfering with decision processes in a current trial (Nakashima and Sugita, 2017; Pavan et al., 2012). Observers were not provided with trial-to-trial feedback but with run-to-run feedback, which was to show the percent correct for a run of 26 trials during each break between scan runs. Before the main fMRI runs, observers inside the MRI scanner repeated what they did during the training sessions by performing 54 practice trials and then 180 threshold-calibration trials. While performing the threshold-calibration trials, which were apart by 2.7s, observers received trial-to-trial feedback based on the classification criterion with radius of 2.84°. A Weibull function was fit to the psychometric curves obtained from the threshold-calibration trials using a maximum-likelihood procedure. From the fitted Weibull function, the size threshold (i.e., the size difference between M ring and L or S ring) associated with 70.7% correct proportion was estimated. We expected that these calibration and practice trials, in addition to the repetitive alternations between the trial-to-trial-feedback calibration trials and the run-to-run-feedback practice trials during the training sessions (see above), would help observers perform the task during the main session in the same way as they did with trial-to-trial feedback. This expectation was supported by the results that the performance levels in the main session (proportion correct = 75±7%) were similar to those targeted by the threshold-estimation procedure (70.7%).

### Behavioral data acquisition for DS#2~4

DS#2-4 were borrowed from the works conducted with different purposes in our lab (Table 1). The specifics of these data sets were as follows: DS#2 was previously published in our previous studies (Choe et al., 2014, 2016) and contributed by 23 observers, each of whom performed 162 trials on the ring stitmuli of 3 different sizes (2.80° ~ 2.88°), with run-to-run and deterministic feedback (see below for definition of deterministic feedback) and inter-trial interval of 13.2s. DS#3 was collected from the same participants of DS#2 but not published before. Each of the subjects, except for one, performed 630 trials on the ring stimuli of 16 different ring sizes (2.72° ~ 2.95°) presented via a staircase method (1-up-2-down), with trial-to-trial and deterministic feedback and inter-trial interval of 2s. The one exceptional subject performed 315 trials. DS#4 was from our preliminary poster presentation (Lee et al., 2016) and contributed by 30 observers, each of whom performed 1,700 trials on the ring stimuli of 5 different ring sizes (3.84°, 3.92°, 4.00°, 4.08 °, 4.16°), with trial-to-trial and stochastic feedback (see below for definition of stochastic feedback) and inter-trial interval of 2.5s. The feedback correctness was determined by a constant criterion for the deterministic feedback condition (DS#1-3) or by a noisy criterion (i.e., the criterion was slightly jittered around a mean, with a gaussian distribution, across trials) for the stochastic feedback condition (DS#4). Stimulus duration was 0.3s for all the data sets. Unlike DS#1, DS#2-4 were acquired without any intensive training sessions.

### Imaging data acquisition and preprocessing

MRI data were collected using a 3 Tesla Siemens Tim Trio scanner equipped with a 12-channel Head Matrix coil at the Seoul National University Brain Imaging Center. Stimuli were generated using MATLAB (MathWorks) in conjunction with MGL (http://justingardner.net/mgl) on a Macintosh computer. Observers looked through an angled mirror attached to the head coil to view stimuli displayed via an LCD projector (Canon XEED SX60) onto a back-projection screen at the end of the magnet bore at a viewing distance of 87 cm, yielding a field of view of 22×17°.

For each observer, we acquired three types of MRI images, as follows: (1) 3D, T1-weighted, whole-brain images (MPRAGE; resolution, 1×1×1 mm; field of view (FOV), 256 mm; repetition time (TR), 1.9 s; time for inversion, 700ms; time to echo (TE), 2.36; and flip angle (FA), 9°), (2) 2D, T1-weighted, in-plane images (MPRAGE; resolution, 1.08×1.08×3.3 mm; TR, 1.5s; T1, 700 ms; TE, 2.79ms; and FA, 9°), and (3) 2D, T2*-weighted, functional images (gradient EPI; TR, 2.2s; TE, 40ms; FA, 73°; FOV, 208mm; image matrix, 90×90; slice thickness, 3mm with 10% space gap; slice, 32 oblique transfers slices; bandwidth, 790Hz/ps; and effective voxel size, 3.25×3.25×3.3mm).

For univariate analysis, the images of individual observers were normalized to the MNI template using the following steps: motion correction, coregistration to whole-brain anatomical images via the in-plane images (Nestares and Heeger, 2000), spike elimination, slice timing correction, normalization using the SPM DARTEL Toolbox (Ashburner, 2007) to 3×3×3mm voxel size, and smoothing with 8×8×8mm full-width half-maximum Gaussian kernel. All the procedures were implemented with SPM8 and SPM12 (http://www.fil.ion.ucl.ac.uk.spm) (Friston et al., 1996; Jenkinson et al., 2002), except for spike elimination for which we used the AFNI toolbox (Cox, 1996). The first six frames of each functional scan (the first trial of each run) were discarded to allow hemodynamic responses to reach a steady state. Then, the normalized BOLD time series at each voxel, each run, and each subject were preprocessed using linear detrending and high-pass filtering (132 s cut-off frequency with a Butterworth filter), conversion into percent-change signals, and correction for non-neural nuisance signals by regressing out mean BOLD activity of cerebrospinal fluid (CSF).

To define anatomical masks, probability tissue maps for individual participants were generated from T1-weighted images, normalized to the MNI space, and smoothed as were done for the functional images by using SPM12, and then averaged across participants. Finally, the locations of CSF, white matter, and gray matter were defined as respective groups of voxels in which the probability was more than 0.5.

The preprocessing steps for multivoxel analysis were the same, but only spatial smoothing was omitted to prevent blurring of the pattern of activity. Unfortunately, in a few of the subjects, functional images did not cover entire brain areas. Voxels in which data were derived from fewer than 17 subjects were excluded for further analysis, which included those in the temporal pole, orbitofrontal, and posterior cerebellum.

### NBMC: generative model

The generative model is the observers’ causal account for noisy sensory measurements, where true ring size, Z, causes a noisy sensory measurement on a current trial, m_(t)_, which becomes noisier as trials elapse, thus turning into a noisy retrieved measurement of the value of Z on trial t − i, r_(t−i)_. Hence, the generative model can be specified with the following three probabilistic terms: a prior of Z, p(Z), a likelihood of Z given m_(t)_, p(m_(t)_|Z), and a likelihood of Z given r_(t−i)_, p(r_(t−i)_|Z). These three terms were all modeled as normal distribution functions, the shape of which is specified with mean and standard deviation parameters, μ and σ: μ_0_ and σ_0_ for the prior, 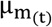 and 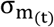 for the likelihood for m_(t)_, and 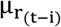 and 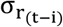 for the likelihood for r_(t−i)_. The mean parameters of the two likelihoods, 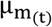 and 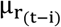, are identical to m_(t)_ and r_(t−i)_; therefore, the parameters that must be learnt are reduced to μ_0_, σ_0_, 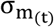, and 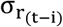. 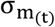 is assumed to be invariant across different values of m_(t)_, as well as across trials. Therefore, 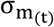 is reduced to a constant σ_m_. Finally, because 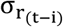 is assumed to originate from σ_m_ and to increase as trials elapse (Gorgoraptis et al., 2011; Zokaei et al., 2015), 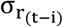 is also reduced to the following parametric function: 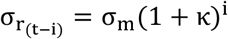, where κ > 0. As a result, the generative model is completely specified by the four parameters, Θ = {μ_0_, σ_0_, σ_m_, κ}.

We defined the ring size, Z, in a normalized space with an arbitrary unit. Specifically, physical ring sizes in visual angle unit were normalized to Z values by dividing their deviances from the M-ring size with the minimum deviance from the M-ring size. For example, five ring sizes in visual degree {1,5,7,9,13} were normalized to Z values of {−3, −1,0,1,3} by dividing the deviances from 7 ({−6, −2,0,2,6}) with the minimum deviance from 7 (2).

### NBMC: stimulus inference (s)

A Bayesian estimate of the value of Z on a current trial, s_(t)_, was defined as the most probable value of a posterior function of a given sensory measurement m_(t)_, which minimizes the mean squared error of estimation for Gaussian distributions (Jaynes, 2003; Körding and Wolpert, 2004). The posterior p(Z|m_(t)_) is a conjugate normal distribution of the prior and likelihood of Z given the evidence m_(t)_, whose mean 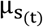 and standard deviation 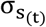 were calculated as follows:

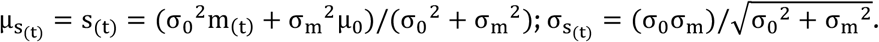

Whenever Z takes one of the ring sizes on each trial as Z_(t)_, its generative noise in generating m_(t)_ was assumed to be equivalent to σ_m_. Therefore, σ_m_ propagates through the Bayesian estimates of stimulus s_(t)_, which results in the sampling distribution of estimates whose mean 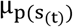 and standard deviation 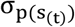 were calculated as follows:

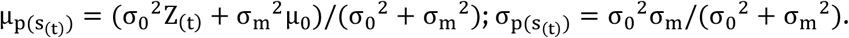

### NBMC: criterion inference (c)

The Bayesian observer estimates the value of criterion on a current trial, c_(t)_, by inferring the most probable value of a posterior function of a given set of retrieved sensory measurements 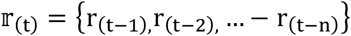, where the maximum number of measurements that can be retrieved, n, was set to 7. This definition of c_(t)_ allows the NBMC to obtain the optimal estimate of c_(t)_ by minimizing squared error (Jaynes, 2003; Körding and Wolpert, 2004). Here, 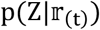 is a conjugate normal distribution of the prior and likelihoods of Z given the evidence 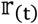, 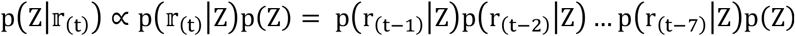, whose mean and standard deviation were calculated (Bromiley, 2003) based on the knowledge of how the retrieved stimulus becomes noisier as trials elapse:

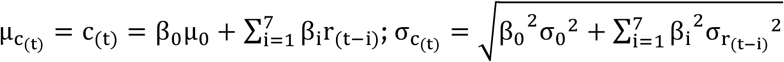

where 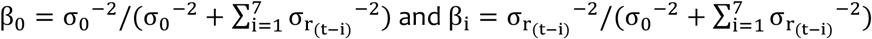.

Much like stimulus estimates, the sampling distribution of criterion estimates have a mean 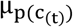 and a standard deviation 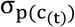 due to generative noise propagation, and they were calculated as follows:

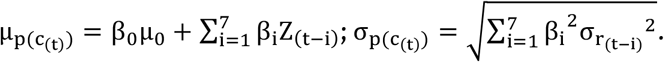

We could analytically derive this equation because we assumed that, when the criterion is inferred, the retrived stimulus measurement r_(t−i)_ is randomly resampled from a normal distribution whose mean is equal to the stimulus that instigated it, Z_(t−i)_, and whose standard deviation is 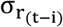. In other words, retrieved sensory measurements ({r_(t−1)_, r_(t−2)_, & − r_(t−n)_}) were assumed to be independent from their initial sensory measurements ({m_(t−1)_, m_(t−2)_, & − m_(t−n)_}) and from one another as well.

### NBMC: deduction of decision variable (v), decision (d) and decision uncertainty (u)

On each trial, the Bayesian observer makes a binary decision d_(t)_ by comparing s_(t)_ and c_(t)_, such that d_(t)_ = ‘large’ if s_(t)_ > c_(t)_, and d_(t)_ = ‘small’ if s_(t)_ < c_(t)_. Thus, given the generative noise propagation, the fraction of ‘large’ choices, pL_(t)_, was defined as the proportion of the bivariate sampling distribution that satisfied the inequality of 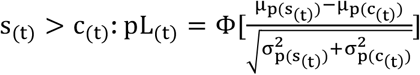 where Θ is cumulative normal distribution (Marzban, 2004).

However, decisions are not made deterministically but probabilistically, because s_(t)_ and c_(t)_ have their respective imprecisions parameterized with 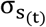 and 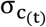. Thus, when committing to a decision with s_(t)_ and c_(t)_, the Bayesian observer calculates the probability of s_(t)_ > c_(t)_, which is called the decision variable, v_(t)_, defined as 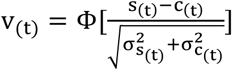, and the decision uncertainty, u_(t)_, which represents the odds that the current decision will be incorrect (Sanders et al., 2016), as follows: 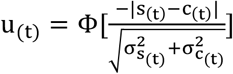.

### Fitting the parameters of NBMC

For each human observer, the parameters of the generative model, 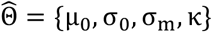, were estimated as those maximizing the sum of log likelihoods for T individual choices made by the observer, 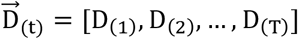:

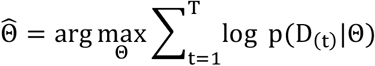

where p(D_(t)_|Θ)=pL_(t)_ for D_(t)_=‘large’ and p(D_(t)_|Θ)= 1 − pL_(t)_ for D_(t)_=‘smalll’.

For each subject, estimation was carried out in the following steps: First, we found local minima for parameters using a MATLAB function, ‘fminseachbnd.m’, with the iterative evaluation number set to 50. We repeated this step by choosing 1,000 different initial parameter sets, which were randomly sampled within uniform prior bounds, and acquired 1,000 candidate sets of parameter estimates. Second, from these candidate sets of parameters, we selected the top 20 in terms of goodness-of-fit (sum of log likelihoods) and searched the minima using each of those 20 sets as initial parameters by increasing the iterative evaluation number to 100,000 and setting tolerances of function and parameters to 10^−7^ for reliable estimation. Finally, using the parameters fitted via the second step, we repeated the second step one more time. Then, we selected the parameter set that showed the largest sum of likelihoods as the final parameter estimates.

We discarded the first trial of each run, the trials in which RTs were too short (less than 0.3 s), and the trials in which no responses were given for parameter estimation for any further analyses. We discarded the first trial of each run for two reasons: first, the criterion could not be inferred on the first trial since there is no measurement to be retrieved yet; second, the first trial is susceptible to non-specific fMRI signals that are irrelevant to the task. Note that some aspects of the fitting procedure are arbitrary (e.g., the parameter boundaries and the number of initial parameter randomization). To be sure, the outcomes of the fitted model parameters are subect to slight changes depending on how those aspects are arbitrated. However, we confirmed that such changes are sufficiently small (results were not shown here) not to affect the main claims of the current study. The fitting codes and data are available in GitHub (see Methods: Data and code accessibility).

### The constant-criterion model

The constant-criterion model has two parameters, bias of classification criterion μ_0_ and measurement noise σ_m_. Stimulus estimates, s_(t)_, were assumed to be sampled from a normal distribution, 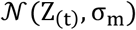. Each stimulus sample has uncertainty 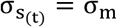. Criterion estimate c_(t)_ was assumed to be a constant, μ_0_; thus 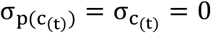.

### Comparing the Bayesian and human observers in decision behavior

Estimating the model parameters separately for each human observer 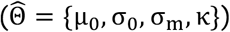 allowed us to create the same number of Bayesian observers, each tailored to each human individual. By repeating the experiment on these Bayesian observers using the stimulus sequences that were identical to those presented to their human partners, we acquired a sufficient number (10^6^ repetitions) of simulated choices, d_(t)_, and decision uncertainty values, u_(t)_, which were determined by the corresponding number of stimulus estimate, s_(t)_, and criterion estimate, c_(t)_, that were randomly sampled from the stimulus (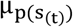 with 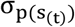) and criterion (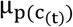 with 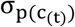) estimate distributions, respectively, on each trial for each observer. Then, the averages across those 10^6^ simulations were taken as the final outcomes. When predicting s_(t)_, c_(t)_, v_(t)_, and u_(t)_ for the observed choice D_(t)_, we only counted the simulation outcomes in which the relation between s_(t)_ and c_(t)_ (i.e., d_(t)_) matched the observed choice D_(t)_.

To check the correspondence in the impact of previous stimuli and previous choices on current choices, we logistically regressed the human current choices onto stimuli and choices using the following model to obtain regression coefficients 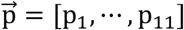 for each observer: 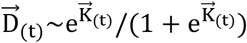, where 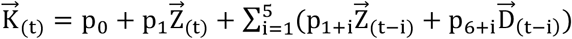, the independent variables were each standardized into z-scores for each observer, and 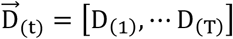, 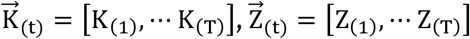, where T is the number of trials in a session. The Bayesian observers’ choices were also regressed with the logistic regression model by substituting 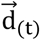 for 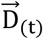, where 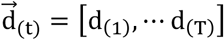. The regression was repeatedly carried out for each simulation, and the coefficients 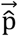 that were averaged across simulations were taken as final outcomes. Due to the additional time consumption associated with regression, the number of simulations for regression was compromised to 10^5^ repetitions, which is smaller than that for pL_(t)_ prediction (10^6^). However, we confirmed that the simulation number was sufficiently large to produce stable simulation outcomes.

When checking the correspondence between the decision uncertainty values u_(t)_ of the Bayesain observers and the RT measurements of the human observers, we converted u_(t)_ values into the values that are comparable to RT measurements by regressing RT measurements onto u_(t)_ using a generalized linear mixed model (GLMM), 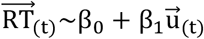, with a random effect of individual observers, where 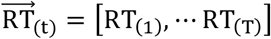 and 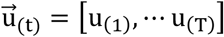. Then, ‘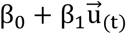’ were pitted against the RT measurents.

To check the correspondence in the impact of previous stimuli on decision uncertainty, we regressed the decision uncertainty estimates 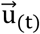 of the Bayesian observers and the RTs of the human observers 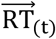, respectively, on current trials onto the congruency of current and past stimuli with a current choice using the following GLMM with a random effect of individual observers: 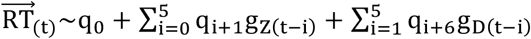, where g_Z(t−i)_ = Z_(t−i)_ × D_(t)_; g_D(t−i)_ = D_(t−i)_ × D_(t)_; D_(t)_ = −1 and 1 for ‘large’ and ‘small’ decisions respectively; for the definition of Z_(t)_ see NBMC: generative model. Here, the independent and dependent variables were both standardized into z-scores for each observer. For comparison with the regression coefficients for the human observers, the regression coefficients for the Bayesian observers were calculated as 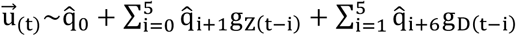by GLMM with a random effect of individual observers, and then linearly re-scaled to those for the human observers using the following GLM: 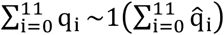, where γ is a GLM parameter for scaling.

### Regression of fMRI responses onto model estimates of decision uncertainty

To identify the brain regions wherein fMRI responses were correlated with u_(t)_ estimates of the NBMC, we regressed the fMRI responses, B_(v,t,i)_, of each voxel, v, at each of the six time points, i, within a single trial, t, onto u_(t)_ using the following GLMM with random effects of individual observers: 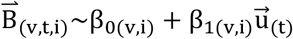, where 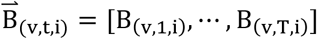. The independent and dependent variables of the regression were both standardized into z-scores for each observer across trials. We first localized the cortical sites where β_1(v,i)_ were statistically significant after correcting for the false discovery rate (FDR) (Benjamini and Hochberg, 1995) over the entire brain voxels tested at each time point, i. Then, the regions of interest (ROIs) were defined as the voxel clusters that cover a region larger than 350 mm^3^ (> 12 contiguous voxels) in which the voxels’ FDR-corrected p-values were less than 0.05 and raw p-values were less than 10^−4^. For ROI analysis, fMRI responses were averaged across individual voxels within a given ROI, and their correspondences with the model estimates of decision uncertainty were calculated using the same procedure as that used for the analysis of RT data.

### Bayes factor analysis for statistical judgment of distinguishable and indistinguishable classifications

Because the NBMC specifies each and every trial with a set of expected values of {s_(t)_, c_(t)_, pL_(t)_, u_(t)_}, it can deterministically predict which trial pairs (i, j) are distinguishable or indistinguishable in decision probability (ΔpL_(i,j)_ = pL_(i)_ − pL_(j)_) and decision uncertainty (Δu_(i,j)_ = u_(i)_ − u_(j)_) based on the metameric currency for each trial pair ([s_(i)_ − c_(i)_] − [s_(j)_ − c_(j)_]).

To statistically judge whether a given trial pair is significantly distinguishable or indistinguishable, we applied the following procedure on DS#1. First, we grouped individual trials according to physical ring sizes, and sorted the trials within each stimulus group by 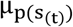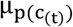 into 10 equal-sized bins, resulting in a total of 30 bins. Second, we made all possible bin pairs out of these bins (900 bin pairs = 30 bin × 30 bins). Then, for each bin pair (i,j), we judged whether the observed ΔpL_(i,j)_ is significantly “equal to zero (H_I_: ΔpL_(i,j)_ = 0)” or “not equal to zero (H_D_: ΔpL_(i,j)_ ≠ i) by calculating the Bayes factor B (Dienes, 2008, 2014; Good, 1979; Jeffreys, 1961; Kass and Raftery, 1995; Wod, 1985), which represents the ratio of the marginal likelihoods of two competing hypotheses, H_I_ (“Indistinguishable”) and H_D_ (“Distinguishable”):

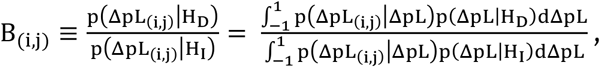

where, p(ΔpL|H_I_) and p(ΔpL|H_D_) represent how likely H_I_ and H_D_, respectively, are for a given value of ΔpL, and p(ΔpL_(i,j)_|ΔpL) represents how likely ΔpL is for a given value ΔpL_(i,j)_. These three likelihoods were assumed to be normally distributed, their means (*μs*) and standard deviation (*σs*) were estimated as follows: *μs* of p(ΔpL_(i,j)_|H_I_) and p(ΔpL_(i,j)_|H_D_) were both zero; *σs* of p(ΔpL_(i,j)_|H_I_) and p(ΔpL_(i,j)_|H_D_) were estimated by computing the SD of ΔpL_(i,j)s_ for the 30 bin pairs where i = j (i.e., SD = 0) and for the 870 bin pairs where i ≠ j; for p(ΔpL_(i,j)_|ΔpL), *μ* was ΔpL_(i,j)_, and *σ* was estimated by computing the across-subject standard error of ΔpL_(i,j)_ for each bin pairs (thus, *σ* slightly differed across the bin pairs). Lastly, for each bin pair, the observed ΔpL_(i,j)_ was judged to be significantly equal to zero and different from zero when the Bayes factor was smaller than 1/3 and greater than 3, respectively, following the convention (Jeffreys, 1961).

The almost identical analyses were applied to the RT and dACC data, except that the metameric currency for u_(t)_ was defined in an absolute term, |s_(t)_ − c_(t)_|. Therefore, two trials become metamers when they have sufficiently similar values of |s_(t)_ − c_(t)_| despite their difference in stimulus-decision congruency (u_(i)_ ≈ u_(j)_|Cong._(i)_ ≠ i)g. i_(j)_) or anti-metamers when they have sufficiently different values of |s_(t)_ − c_(t)_|, despite their similarity in stimulus-decision congruency (u_(i)_ ≠ u_(j)_|Cong._(i)_ = Cong._(j)_).

### Searching for multivoxel patterns of activity signaling the latent variables of the NBMC

To decode the latent variables of the NBMC in fMRI responses, the time-resolved support vector regression (SVR) was carried out in conjunction with a searchlight technique (Haynes, 2015; Kahnt et al., 2011b). At each of the first four time points, i, within a single trial, t, B(e,k,t,i) (which represents the preprocessed but unsmoothed fMRI responses of a voxel cluster centered at a gray-matter voxel, e, where k where × denotes an entire set of voxels that comprise the cluster) was selected as a searchlight. The first four time points were chosen given that the latent variables must precede u_(t)_, which was detected at the fourth time point of the univariate fMRI responses in the dACC. Although the searchlight cluster had a radius of 9mm and thus consisted of 123 voxels, the exact number of voxels in each searchlight varied, because the voxels located in CSF or white matter were discarded as non-neural signals. For each observer, the latent variables (s_(t)_, c_(t)_, and v_(t)_) were decoded for each searchlight using the cross-validation method of one-run-leave-out (8-fold cross-validation).

We performed the SVR using the LIBSVM (http://www.csie.ntu.edu.tx/~sjlin/libsvm) with a linear kernel and constant regularization parameter of 1. For each voxel and each subject, B_(e,k,t,i)_ and the values of the latent variables were z-scored across trials before decoding. To calculate the significance level of the decoded information, each of the 3D spatial maps of the decoded values of the latent variables (v_B(e,t,i)_, c_B(e,t,i)_, or s_B(e,t,i)_) was smoothed with a 5 mm FWHM Gaussian kernel. We then regressed the smoothed v_B(e,t,i)_, c_B(e,t,i)_, and s_B(e,t,i)_ onto v_(t)_, c_(t)_, and s_(t)_, respectively, using 16, 13, and 13 regression models, respectively. Those regression models were deduced from the causal structure between the latent variables of the NBMC (see the next section). We concluded that a given cluster carries the neural signals of v_(t)_, c_(t)_, or s_(t)_, only when all those regression models are satisfied over 12 contiguous searchlights. For the ROI analysis, the decoded values of a given latent variable were averaged over the all searchlights within each ROI. The brain imaging results were visualized using xjView toolbox for the cross sectional images and Connectome Workbench (Marcus et al., 2011) for the inflated images.

### Deduction of the regression models for v_(t)_, c_(t)_, and s_(t)_

To identify brain signals of c_(t)_, s_(t)_, and v_(t)_, we defined three a priori lists of regressions that must be satisfied by the brain signals. We stress that each of these lists consists of the necessary conditions to be satisfied, and the conditions that must not be satisfied as well, because both significant (βof the ne) and non-significant (β a) regressions are deduced from the causal structure of the variables in the NBMC. The lists were as follows.

The 13 regressions for the brain signal of c: (c1-3), y_c_, c decoded from brain signals, must be regressed onto c—the variable it represents—even when c is orthogonalized to v or d, because it should reflect the variance irreducible to the offspring variables of c; (c4), y_c_ must not be regressed onto s because c and s are independent; (c5,6), y_c_ must be regressed onto v but not when v is orthogonalized to c because the influence of c on v is removed; (c7,8) y_c_ must be regressed onto d but not onto u because u cannot be correlated with c unless d or s is fixed; (c9-11), y_c_ must be regressed onto, not the current stimulus, but the past stimuli—strongly onto the 1-back stimulus and more weakly onto the 2-back stimulus (thus, non-significant regression with one-tailed regression in the opposite sign is modeled conservatively); (c12,13), y_c_ must not be regressed onto previous decisions at all because c is inferred solely from retrieved stimulus measurements.

The 13 regressions for the brain signal of s: (s1-3), y_s_, s decoded from brain signals, must be regressed onto s—the variable it represents per se—even when s is orthogonalized to v or d because it should reflect the variance irreducible to the offspring variables of s; (s4), y_s_ must not be regressed onto c because s and c are independent of each other; (s5,6), y_s_ must be regressed onto v but not when v is orthogonalized to s because the influence of s on v is removed; (s7,8) y_s_ must be regressed onto d but not onto u because u cannot be correlated with s unless d or c is fixed (s9-11), y_s_ must be regressed onto the current stimuli and not the past stimuli because s is inferred solely from the current stimulus measurement; (s12,13), y_s_ must not be regressed onto previous decisions because s is inferred solely from the current stimulus measurement.

The 16 regressions for the brain signal of v. (v1-4), y_v_, v decoded from brain signals, must be regressed onto v—the variable it represents—even when v is orthogonalized to c, s, or d, because it should reflect the variance irreducible to the offspring variables of v; (v5,6), y_v_ must be regressed onto one of its parents c, but not when c is orthogonalized to v, because the influence of c on v is removed; (v7,8), y_v_ must be regressed onto one of another parent s, but not when s is orthogonalized to v, because the influence of s on v is removed; (v9,10), y_v_ must be regressed onto d but not onto u because u’s correlation with its parent v cannot be revealed without holding the variability of d; (v11-13), y_v_ must be positively regressed onto the current stimulus because the influence of the current stimulus on v is propagated via s, and negatively regressed onto the past stimuli because the influence of the past stimuli on v is propagated via c—strongly onto the 1-back stimulus and more weakly onto the 2-back stimulus (thus, non-significant regression with one-tailed regression in the opposite sign is modeled moderately); (v14-16), y_v_ must be regressed onto the current decision and not the past decisions because the current decision is a dichotomous translation of v, whereas past decisions have nothing to do with the current state of v.

### Bayesian network analysis

For the data-driven Bayesian network analysis, we derived an exhaustive set of causal structures between the brain signals of the latent variables of the NBMC and calculated BIC for each structure (Scutari, 2009). Specifically, to see whether the c, s and v signals are consistent with the causal graph structure that is prescribed a priori by the NBMC, we searched for the causal graph (G) whose likelihood is maximal given the time series of three brain signals, one for each of the latent variables 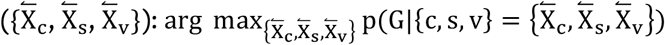. A total of 27 causal graphs can be created out of three variables nodes (3 edges × 3 edges × 3 edges between the nodes); a total of 6 triplets of brain signals can be used for 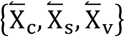 since we have three (IPL_c1_, STG_c3_, STG_c5_), two (DLPFC_s3_, Cereb_s5_), and single (STG_v5_) candidate brain signals for c, s, and v, evaluated which of the 162 (27 edges structures × 6 nodes combinations) possible Gs is most likely. Since these Gs differ in complexity, i.e., the number of parameters, we used the Bayesian Information Criterion (BIC) values for comparison (the smaller a BIC value is, the more likely a graph is).

### The categorization model

To demonstrate the history effects in the categorization model, we adapted the “exponentially weighted moving average” model that was developed by Norton and her colleagues to the sequences of stimuli in DS#1 (see Norton et al., 2017 for model description). The likelihoods of two categories (‘small’ and ‘large’ categories) over ring sizes were assumed to be gaussian distributions. A model parameter σ_m_ determined the trial-to-trial noise of sensory measurement, m_(t)_. As σ_m_ becomes greater, a sensory measurement becomes noisier (more deviated from the true stimulus). The model used the arithmetic mean of the two category likelihoods 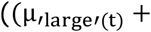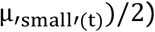, where the two category likelihoods are equal, as the decision boundary, such that ‘small’ and ‘large’ decisions were made when a sensory measurement m_(t)_ was smaller and larger, respectively, than the boundary. After the trial t, the model updates only the mean of likelihood for one category that corresponds with a past decision by shifting the old mean toward a past stimulus with an exponential decaying weight, which is determined by a parameter, α. Note that trial-to-trial feedback determined the category membership in the orginal study whereas a decision did so in the current study since there was no trial-to-trial feedback in DS#1. For example, when a ‘large’ decision has been made, only the mean of the ‘large’ category is updated such that 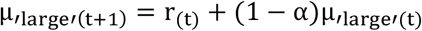 while the mean of the ‘small’ category remains unchanged 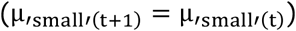, where r_(t)_ is a reinstated sensory measurement, the noisiness of which is also governed by σ_m_. As α becomes greater, the mean of the category likelihood becomes more attracted toward the past stimulus.

### Statistical tests

To calculate confidence intervals, a set of bootstrap-sampled data was obtained by resampling 10^5^ times with repetition, and the mean and interval size of the threshold (e.g., 95%) were then computed for the bootstrap data set. For all statistics of DS#1, 18 individuals were used except the whole brain analysis, in which statistics at some regions of the ventral lobe were calculated with 17 individuals. For DS#2-4, the numbers of subjects for statistical test are provided in Table 1. Statistical significances were calculated using two-tailed tests, except for the regression models for decoding the latent variables of the NBMC from fMRI responses.

### Data and code accessibility

Codes and data for fitting NBMC to behavior data can be found in https://github.com/Heeseung-Lee/LeeLeeChoeLee2020_JN.

## Results

### Core properties of classification

To gain intuitions about the core properties of classification that the NBMC aims to capture, consider the scenario depicted in Figure 2. There are three apple farms that differ in fertility (‘B(arren)’, ‘O(rdinary)’, and ‘F(ertile)’), and thus the overall apple size tends to be small, medium, and large in the B, O, and F farms, respectively. Farmers’ job is to pick apples and sort them in size into ‘small’ and ‘large.’ In the first year, three farmers were all working in the same O farm (Fig. 2*A*, left side). In the second year, one farmer continued to work in O farm, but the remaining two moved to B and F farms respectively (Fig. 2*A*, right side). Since the apple-sorting criterion for a given farmer reflects the typical size of the apples in the farm, we can readily intuit that the criteria were similar among the farmers in the first year but have become different in the second year. This intuition relates to two historical properties of classification: (P1) c varies as a function of past experiences of stimulus, and (P2) c is more strongly attracted toward recent experiences than old ones.

**Figure 2.**
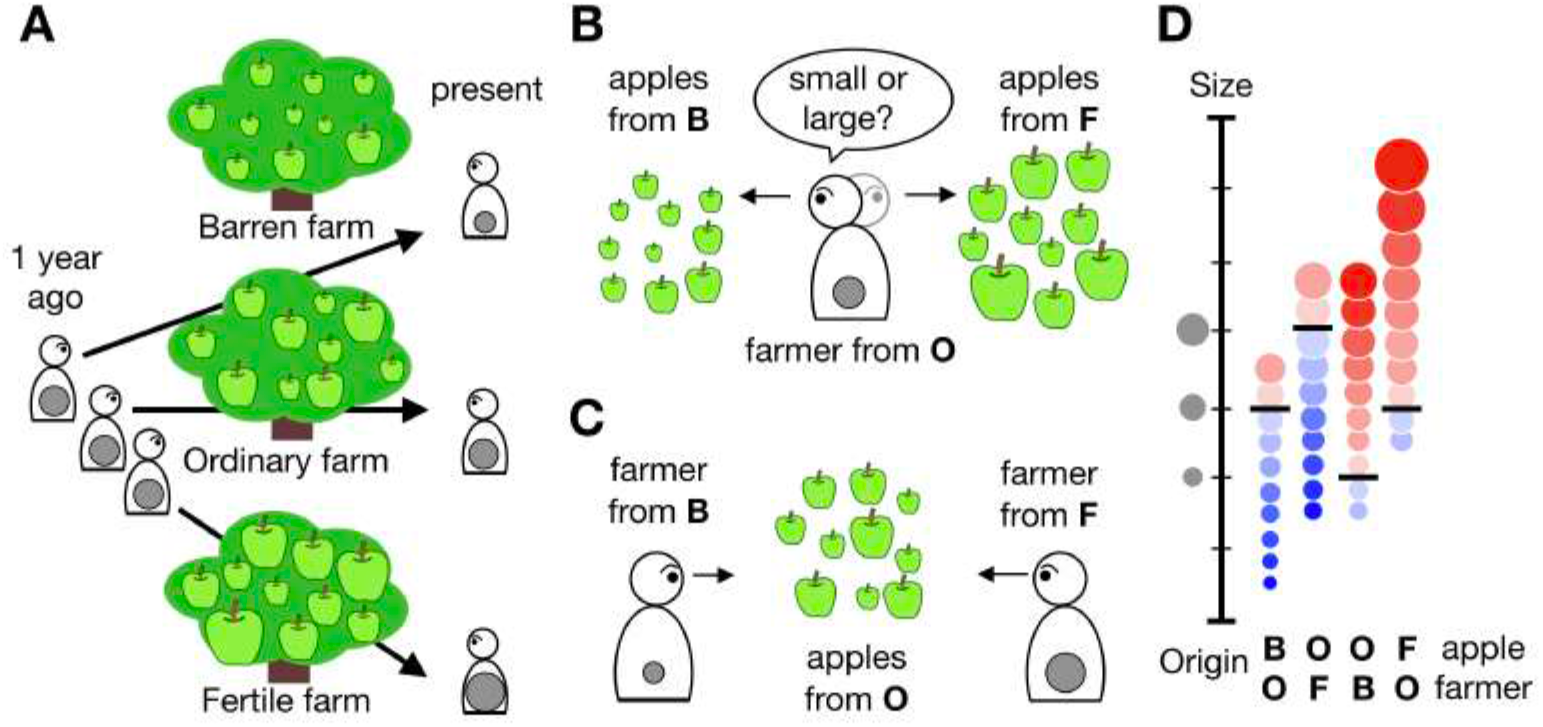
The formation, use, and consequences of a criterion in an apple-classification scenario. ***A,*** Experience-dependent formation of criteria. In the first year, three farmers all worked at Ordinary farm and classified the apples there into ‘large’ or ‘small’ sizes, forming a criterion that is close to the typical size of the apples of Ordinary farm (gray circles inside the bodies in the left side). In the second year, two of the farmers moved to Barren and Fertile farms respectively. As the farms differ in fertility and thus as their apples differ in size, the three farmers experienced apples of different sizes and form different criteria (gray circles inside the bodies in the right side). ***B,*** Example cases where the apples from different farms are sorted by the same farmer. The farmer from the Ordinary farm classifies the apples from the Barren and Fertile farms as mostly ‘small’ and ‘large’, respectively, by comparing the apple sizes against the medium apple size of Ordinary farm, the criterion. ***C,*** Example cases where the apples from the same farm are sorted by different farmers. Due to the different classification criteria, apples from the Ordinary farm are mostly judged as ‘large’ by the farmer from the Barren farm and ‘small’ by the farmer from the Fertile farm. ***D,*** Distributions of decisions and uncertainties for the four cases shown in *B* and *C*. The black horizontal bars represent a criterion, and the size of circles represents apple size. The hue and saturation of circles represent decision identity (blue for ‘small’, red for ‘large’) and uncertainty (increasing desaturation with increasing uncertainty), respectively.

Let us say that, one day in the second year, the farmers visit one another’s farms and classify the apples there. First, imagine that the farmer from O farm sorts the apples from B farm (‘O sorting B’) or the apples from F farm (‘O sorting F’; Fig. 2*B*, the first and the fourth columns in Fig. 2*D*). We can intuit that farmer O tends to classify apples as ‘small’ in B farm and ‘large’ in F farm, because apple sizes mostly fall below and above the farmer’s criterion, respectively. We can also intuit that the farmer’s decision uncertainty also varies from apple to apple, becoming increasingly uncertain as the apple size falls closer to the classification criterion. This exemplifies the contribution of stimulus to decision variable, when c is fixed. Next, imagine that apples from O farm are sorted by the farmer from F farm (‘F sorting O’) or by the farmer from B farm (‘B sorting O’; Fig. 2*C*, the second and the third columns in Fig. 2*D*). Here we can intuit the contribution of c to decision variable: the very same apples will be sorted differently, and with different degrees of uncertainty, depending on which farmer sorts them.

Intriguingly, different pairings bespeak the relativity of classification in which decision and its uncertainty are determined not by an absolute quantity of stimulus but by the relative quantity of stimulus to c. For example, the ‘O sorting B’ and ‘F sorting O’ cases differ both in absolute apple size and in farmer, but are similar in decision fraction and uncertainty distribution (the first and the second columns in Fig. 2*D*). This is because the differences in stimulus are compensated by the differences in c. Put together, the intuitions earned from the farmers’ cross-visit day relate to the two remaining properties of classification: (P3) s and c have respective contributions to the variability of the decision variable for classification; (P4) interplays between s and c realize the relativity of classification.

### Ring-size classification task

Over consecutive trials, observers sorted rings by size into two classes, ‘small’ and ‘large’, under moderate time pressure (Fig. 1*A*). For the main data set, in which behavioral and fMRI responses were concurrently acquired (DS#1, Table 1), rings of three marginally different radii were used to ensure that decisions were made with uncertainty. Observers were not informed about the actual number of ring sizes. The threshold (70.7% correct) ring sizes were tailored for individuals (see Methods for details). Considering the possibility that ‘correct’ and ‘incorrect’ feedback events evoke the brain activities associated with rewards (Carlson et al., 2011; Marco-Pallarés et al., 2007) or errors (Carter et al., 1998; Cavanagh and Frank, 2014; Holroyd et al., 2004), we did not provide observers with trial-to-trial feedback during the fMRI runs. Instead, we provided observers with run-to-run feedback by showing the proportion of correct responses at the end of each run of 26 trials. The actual level of task performance (mean=75.7%, SD=6.3%) was close to that aimed by the training sessions, which suggests that a negligible amount of perceptual learning occurred during data collection.

The contributions of current stimuli to classification were summarized by plotting the proportion of ‘large’ decisions (pL) and response time (RT) against ring size (Fig. 1*B,C*). As the ring size increased, the pL increased (logistic regression of decisions onto ring size, β = 0.99, P = 6.6×10^−4^), the RT of ‘large’ decisions decreased (linear regression of RT onto ring size, β = −0.17, P = 8.8×10^−13^), and the RT of ‘small’ decisions increased (linear regression of RT onto ring size, β = 0.19, P = 2.4×10^−16^).

When acquiring the main data, we had to adopt specific experimental procedures, including the run-to-run feedback (instead of trial-to-trial feedback), the small (3) number of ring sizes, and the long (13.2 s) inter-trial interval, due to the concurrent collection of behavioral and fMRI data. To show that the validity of the NBMC is not limited to these procedures, we also included additional data sets (DS#2-4; Table 1; (Choe et al., 2014, 2016; Lee et al., 2016)), which were collected outside the MRI scanner with procedures different from those of the main data sets.

### The normative Bayesian model of classification (NBMC)

The generative model, an observer’s account for how their sensory measurements are generated, was assumed as follows: on a trial t, the true size Z_(t)_ is randomly sampled from a prior distribution p(Z), and the observer’s sensory measurement m_(t)_ is another noisy sample from a likelihood distribution p(m|Z) centered around Z_(t)_ (Fig. 3*A*). The size classification task, splitting population size values into small and large halves, can be defined as judging whether Z_(t)_ is larger or smaller than 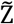, the median of the true ring sizes shown over all trials. Because the observer can access only the measured sizes of stimuli encountered so far, the Bayesian solution is to infer Z_(t)_ and 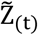, which correspond to inferring a stimulus and a criterion, respectively, from a limited set of noisy measured sizes {m_(1)_, m_(2)_, …, m_(t)_}. We modeled both Z and m as Gaussian random variables, and each was parameterized by the mean and standard deviation (SD).

**Figure 3.**
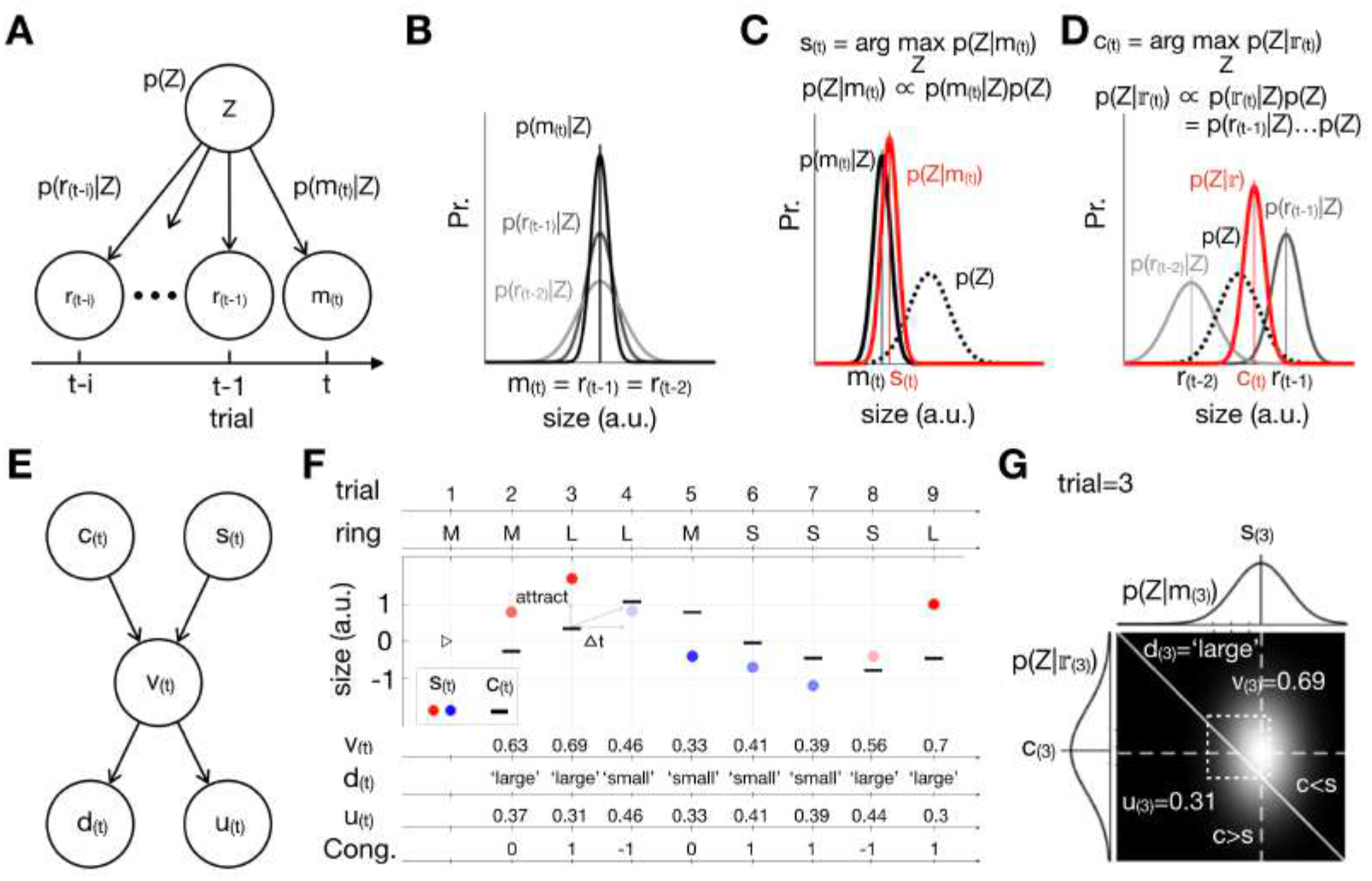
Descriptions of NBMC. ***A,*** Generative model. Arrows indicate conditional dependencies. Z, true stimulus size. m_(t)_, noisy observation on a current trial t. r_(t−i)_, noisy observation retrieved from a past trial t − i. ***B,*** Increases in measurement noise over trials. A likelihood function over ring-size encodes the most likely value of Z_(t)_ and its uncertainty, which increases as trials elapse. ***C,*** Bayesian inference for a current stimulus, s. A posterior function (red) represents s_(t)_ and its uncertainty. ***D,*** Bayesian inference of a criterion, c. A posterior function (red) represents c_(t)_ and its uncertainty. ***E,*** Causal structure of variables. s_(t)_ and c_(t)_ construct v_(t)_, a continuous decision variable. Then, v_(t)_ determines d_(t)_, binary decision, and u_(t)_, decision uncertainty. ***F,*** Illustration of decision episodes over example trials. On the stimuli presented by experimenters, the Bayesian observer makes decisions by comparing s (colored dots) against c (black bars), which is updated continuously over trials. As s_(t)_ and c_(t)_ are inferred, v_(t)_, d_(t)_ and u_(t)_ are deduced accordingly. ‘Cong.’ indicates a congruence between d_(t)_ and ring size and is defined as d_(t)_×Z_(t)_, where d_(t)_ ∈ {−1,1} for ‘small’ and ‘large’, respectively, and Z_(t)_ ∈ {−1,0,1} for S, M, L rings, respectively. Note that c_(t)_ appears to chase after s_(t−i)_ as indicated by the small arrows. ***G,*** Illustration of the model variables in a bivariate decision space for trial 3 in the panel *F*. The light intensity corresponds to the probability density of the bivariate conjugate of p(Z|m_(3)_) and 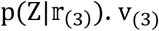 corresponds to the fraction of the bivariate distribution above the identity line, which is greater than 0.5 and thus leads to the ‘large’ decision (d_(3)_ = ′large′). u_(3)_ corresponds to the fraction of the bivariate distribution that is against the decision, i.e., the probability that the decision was incorrect.

Critically, we assumed that there exists an additional source of noise arising from imperfect memory retrieval; the likelihood distribution becomes less reliable as trials elapse (Fig. 3*B*). To distinguish from the likelihood of Z for the current-trial measurement p(m_(t)_|Z), we denoted the likelihood of Z for the measurement ‘retrieved’ from an elapsed trial t − i by p(r_(t−i)_|Z) and assumed that its uncertainty increases as trials elapse (Gorgoraptis et al., 2011; Zokaei et al., 2015), as represented by the widening curves as trials elapse shown in Fig. 3*B*.

On a trial t, the Bayesian observer infers the size of a current stimulus (Z_(t)_) and the criterion 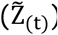 by inversely propagating {…, r_(t−2)_, r_(t−1)_, m_(t)_} over the generative model. The inferred stimulus size, s_(t)_, is the most probable value of Z given the measurement on a current trial, as captured by the following equation:

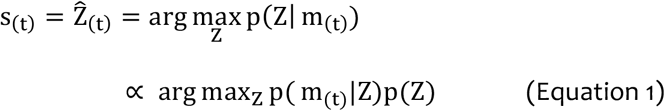

where the variance of the posterior reflects the precision of s_(t)_ (Fig. 3*C*).

On the other hand, the inferred criterion, c_(t)_, which corresponds to the inferred value of 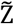, is the most probable value of Z given the measurements over elapsed trials, as captured by the following equation:

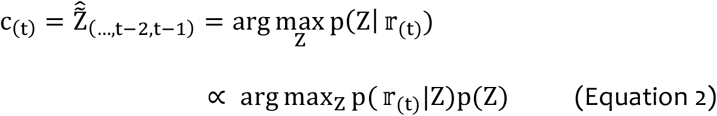

where 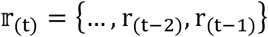. Here, the variance of the posterior, which reflects the precision of c_(t)_, is the optimally weighted sum of variances of the retrieved measurements and the prior size (Fig. 3*D*). Equation 2 implies that c_(t)_ is more attracted towards a recent stimulus than to older ones because the representational uncertainty of working memory increases as trials elapse (Fig. 3*D*). Note that s_(t)_ and c_(t)_ are the optimal inferences of the current stimuli and criterion given the generative model of the Bayesian observer because taking the maximum value of a posterior minimizes estimation errors when the posterior follows a Gaussian distribution (Jaynes, 2003; Körding and Wolpert, 2004).

Having estimated s_(t)_ and c_(t)_, the Bayesian observer performs the task by deducing a decision variable v_(t)_ from s_(t)_ and c_(t)_ and translating it into a binary decision d_(t)_ with a degree of uncertainty u_(t)_ (Fig. 3*E*); v_(t)_ is the probability that s_(t)_ will be greater than c_(t)_ (p(s_(t)_ > c_(t)_)); d_(t)_ is ‘large’ or ‘small’ if v_(t)_ is greater or smaller than 0.5, respectively; u_(t)_ is the probability that d_(t)_ will be incorrect (p(s_(t)_ < c_(t)_|d_(t)_ = ′large′) or p(s_(t)_ > c_(t)_|d_(t)_ = ′small′)) (Sanders et al., 2016).

Figure 3F and 3G graphically summarize how the NBMC works. Figure 3F illustrates how c is updated as a function of past stimuli, and takes part, with s, in classification by determining v and u over an example stream of trials: c tends to run after s, as implied by Equation 2; when s_(t)_ is larger (smaller) than c_(t)_, v_(t)_ was larger (smaller) than 0.5 and thus d_(t)_ = ′large′ (′small′); as s_(t)_ becomes closer to c_(t)_, u_(t)_ becomes higher. Also, note that the congruence between a ring size and d_(t)_ can descriptively capture the dynamic of u_(t)_:u_(t)_ is lower when a stimulus and a decision are incongruent. The bivariate decision space shown in Figure 3G shows a snapshot of how the variables comprising decision outcomes (v_(t)_, d_(t)_, and u_(t)_) are deduced from the posterior probability distributions of s_(t)_ and c_(t)_ on an example (t=3) trial.

In summary, according to the NBMC, as the ring size (Z) varies over trials, the Bayesian observer continues to (i) make the inferences of current stimuli and criteria (s and c), (ii) deduce v from s and c, and (iii) make decisions (d) with varying degrees of uncertainty (u). To verify the implications of the NBMC, we took the following steps. First, for each human observer, we created a Bayesian partner by fitting the NBMC to the decisions of that observer. Second, we made these Bayesian observers simulate decisions (d) and decision uncertainties (u) on the very sequences of stimuli that were encountered by their corresponding human partners. Then, we examined the correspondences between the human and the Bayesian observers regarding each of the aforementioned four core properties of classification (P1~4).

### History effects of past stimuli on classification (P1 and P2)

According to the c-inference algorithm of the NBMC (Equation 2; Fig. 3*D*), c is affected only by past stimuli but not by past decisions, and is attracted toward past stimuli more strongly toward recent ones than toward old ones (Fig. 3*D,F*). Then, according to the deduction algorithm of the NBMC (Fig. 3*E*), such attractive shifts of c toward past stimuli will be translated into the decisions that are repelled away from past stimuli, more strongly from recent ones than from old ones.

This model behavior realizes the two historical properties of classification that were intuited in the apple-classification scenario (P1 and P2).

To check the correspondences between the Bayesian observers and the humans regarding P1 and P2, we regressed current decisions that were made by the Bayesian observers and by the humans onto past stimuli, current stimuli and past decisions (Fig. 4). As implied by the NBMC, for both of the Bayesian observers (Fig. 4*A*) and the humans (Fig. 4*B*), current decisions were more strongly repelled from more recent stimuli (P2), while effects of past decisions on current decisions are negligible (P1). The regression coefficients were well matched between the Bayesian observers and the humans. The above correspondences were also replicated in the other three data sets (Fig. 4*C-E*), which supports the generalizability of the NBMC’s account for the history effects in classification. For all of the data sets, the repulsive history effects of past stimuli on current decisions could not be simulated by an alternative model (‘constant-c model’; Fig. 4*F*), in which c was fixed at a constant instead of being inferred from past sensory measurements (see Methods for detailed model description).

**Figure 4.**
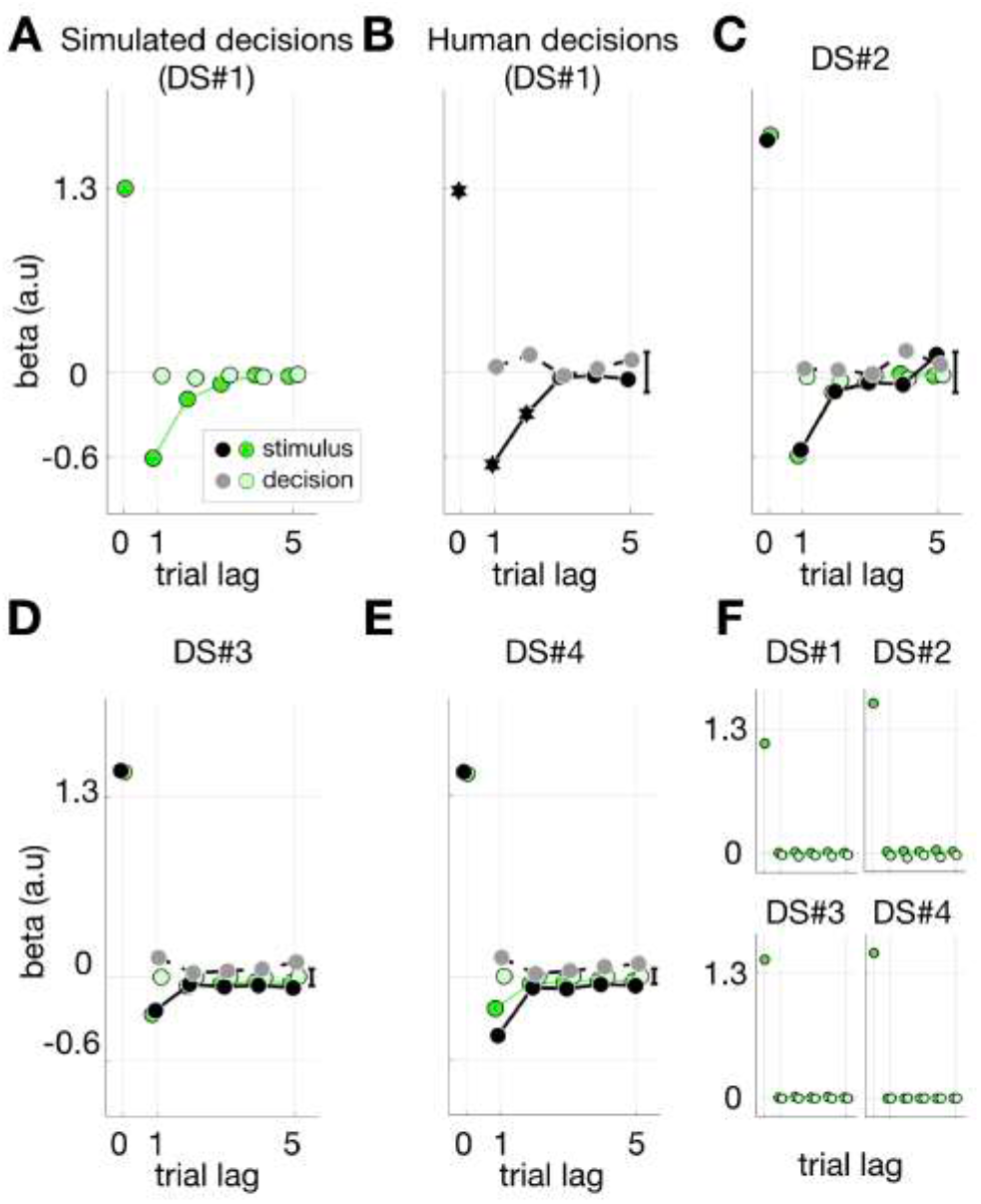
Regression of current decisions onto past stimuli. ***A-B,*** Multiple logistic regressions of the simulated decisions of the NBMC Bayesian observers (A) or the decisions of the human observers (B) onto stimuli and past decisions for DS#1. The regression coefficients are plotted against trial lags, where zero means a current trial. The green and black symbols represent the coefficients for the Bayesian and human observers, respectively. The dark and light symbols represent the coefficients for stimuli and decisions, respectively. For clarity, only the average of 95% bootstrap confidence intervals of the mean coefficients is shown as vertical black bar for the human data. The hexagons indicate the coefficients that are significantly deviated from zero (pairwise t-test, P < 0.05). ***C-E,*** Multiple logistic regressions of the simulated decisions of the NBMC Bayesian observers and the decisions of the human observers onto stimuli and past decisions for DS#2 (C), DS#3 (D), and DS#4 (E), respectively. The color and lightness schemes are identical to those used in A and B. ***F,*** Multiple logistic regressions of the simulated decisions of the constant-c model observers onto stimuli and past decisions for all the data sets. The lightness schemes are identical to those used in A.

The c-inference and deduction algorithms of the NBMC also imply specific history effects of past stimuli on decision uncertainty (u_(t)_): the decision uncertainty on a current trial varies as a function of the congruence of a current decision with only past stimuli, but not with past decisions. This is because, as c_(t)_ is attracted towards a past stimulus, s_(t)_ that would lead to the decision congruent with that past stimulus (e.g., making a ‘large’ decision on a current trial after viewing an L ring on a previous trial) tends to be closer to c_(t)_ and thus be more uncertain compared to s_(t)_ that would lead to the decision incongruent with that past stimulus (e.g., making a ‘small’ decision on a current trial after viewing an L ring in a previous trial). In the case of ‘F sorting O’ in the apple-classification scenario, which is depicted in the second column of Figure 2*D*, the congruent and incongruent conditions correspond to the red and blue circles, respectively. As a result, u_(t)_ is high when d_(t)_ is congruent with past stimuli (P1). And, again because c is more attracted towards recent stimuli than towards old ones, this positive history effect of decision-congruent past stimuli on u_(t)_ should be stronger for recent stimuli than for old ones (P2).

Unlike decisions, decision uncertainty was not directly measured from the human observers. Instead, to compare with the Bayesian observers regarding history effects on decision uncertainty, we used RT as a proxy of decision uncertainty of the human observers based on their tight linkage reported by previous studies (Palmer et al., 2005; Ratcliff and McKoon, 2008; Urai et al., 2017). Then, the simulated values of u_(t)_ and the RT measurements, for the Bayesian observers and the humans respectively, were regressed onto both the congruence of d_(t)_ with past stimuli (‘D-S congruence’) and the congruence of d_(t)_ with past decisions (‘D-D congruence’) (Fig. 5). The patterns of regression coefficients were consistent between the Bayesian observers and the humans while satisfying the history effects on decision uncertainty that are implied by the NBMC. As in the decision data, these results were replicated for all the data sets and could not be simulated with the constant-c model.

**Figure 5.**
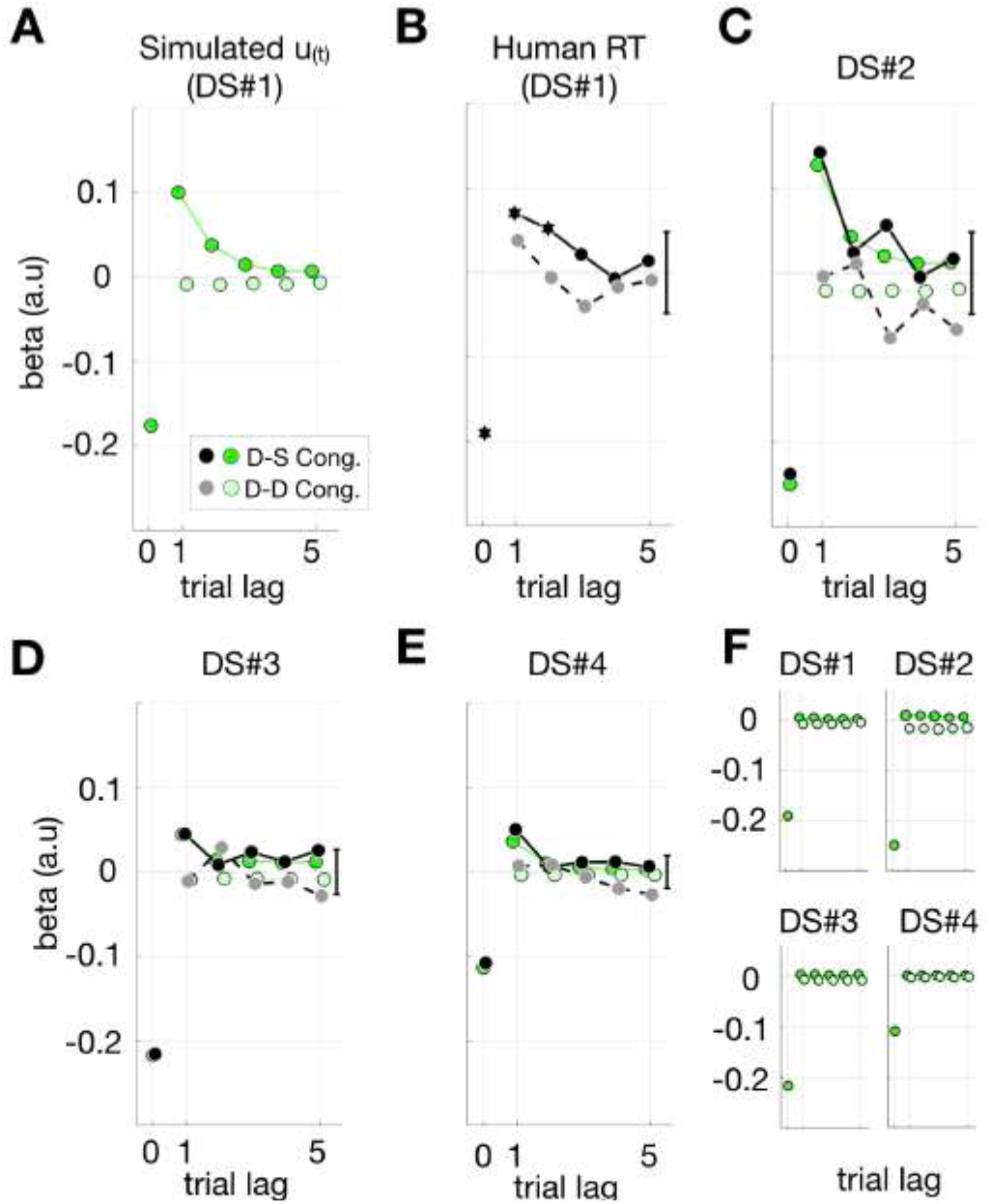
Regression of current decision uncertainty or RT onto the congruence of current decision with past stimuli. ***A-B,*** Multiple linear regressions of the simulated decision uncertainty measures (u_(t)_) of the NBMC Bayesian observers (*A*) or the decision RTs of the human observers (*B*) onto the congruence of current decisions with stimuli or with past decisions for DS#1. The regression coefficients are plotted against trial lags, where zero means a current trial. The green and black symbols represent the coefficients for the Bayesian and human observers, respectively. The dark and light symbols represent the coefficients for the congruence of current decisions with stimuli and past decisions, respectively. For clarity, only the average of 95% bootstrap confidence intervals of the mean coefficients is shown as vertical black bar for the human data. The hexagons indicate the coefficients that are significantly deviated from zero (Generalized multiple linear mixed effect model (GLMM), P < 0.05). ***C-E,*** Multiple linear regressions of the simulated u_(t)_ of the NBMC Bayesian observers and the decision RTs of the human observers onto the congruence of current decisions with stimuli or with past decisions for DS#2 (C), DS#3 (D), and DS#4 (E), respectively. The color and lightness schemes are identical to those used in A and B. ***F,*** Multiple linear regressions of the simulated u_(t)_ of the constant-c model observers onto the congruence of current decisions with stimuli or with past decisions for all the data sets. The lightness schemes are identical to those used in A.

As another means of testing the model implications regarding history effects on decision uncertainty, we examined the fMRI responses from the human observers in search of cortical activities that are correlated with u_(t)_. We found such activities in the dorsal anterior cingulate cortex (dACC) and bilateral anterior insula (Ins), which were known to be associated with decision uncertainty in the literature (Grinband et al., 2006; Shenhav et al., 2014; Sheth et al., 2012). In these regions, the fMRI responses at 5.5s from stimulus onset, which are likely to reflect the neural activities at around the moment of decision making considering the typical hemodynamic delay of fMRI signal, were significantly correlated with u_(t)_ (Table 2; Fig. 6*A,C*).

**Table 2.** Specifications of the brain clusters in which fMRI responses were correlated with decision uncertainty. When a set of constraints (uncorrected P < 10^−4^, FDR-corrected P < 0.05, cluster size > 324 mm^3^ (or contiguous voxels > 12)) was applied, three regions of interest (ROIs) were significantly correlated with u_(t)_ at the fourth fMRI time point (5.5s after stimulus onset). The two-tailed tests were applied for evaluating the significance of GLMM.

| Cortical<br>area | Correlated | Volume | Peak voxel |  |  |
| --- | --- | --- | --- | --- | --- |
|  | time point | (mm<sup>3</sup>) | MNI | GLMM | GLMM |
|  | (s) |  | coordinate | p-value | coefficient |
| | | | | of $u_{(t)}$ | of $u_{(t)}$ |
| Dorsal<br>anterior<br>cingulate<br>cortex | 5.5 | 3564 | [3, 27, 36] | $1.43 \times 10^{-8}$ | 0.095 |
| Left<br>anterior<br>insula | 5.5 | 2268 | [-27, 24, 6] | $4.6 \times 10^{-8}$ | 0.095 |
| Right<br>anterior<br>insula | 5.5 | 2619 | [33, 18, 6] | $8.3 \times 10^{-7}$ | 0.082 |

**Figure 6.**
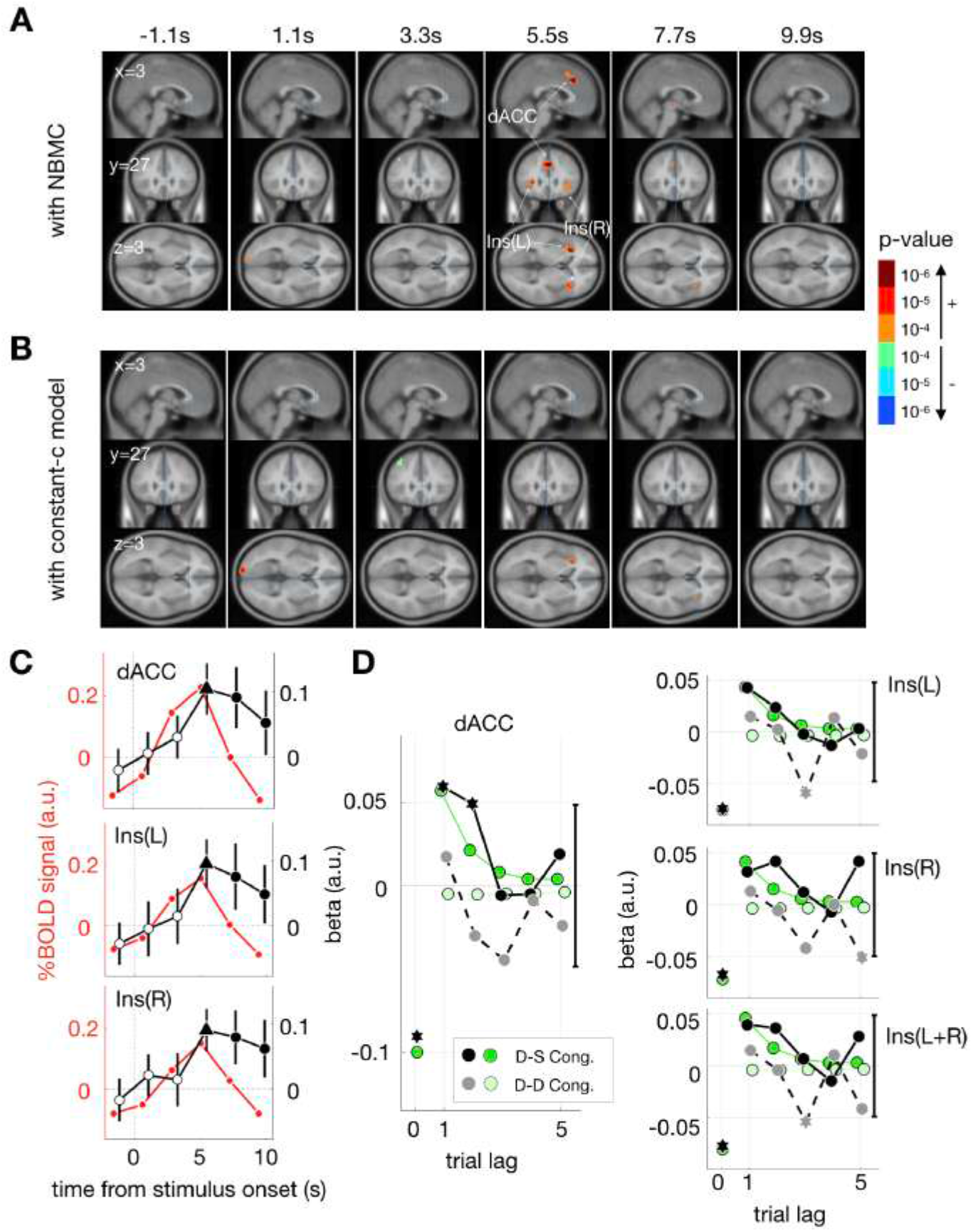
Univariate fMRI responses correlated with decision uncertainty. ***A-B,*** Regressions of fMRI responses at individual voxels onto u_(t)_ for each of the 6 time points that comprise single trials (the numbers at the top indicate the time points from stimulus onset). For each of those time points, the cortical sites where the p-values of regression coefficients were smaller than a threshold (GLMM, P < 10^−4^, uncorrected) were marked in color and overlaid on the sagittal (x=3), coronal (y=27), and axial (z=3) slices of a template brain. ***A,*** Regression results with u_(t)_ of the NBMC. ***B,*** Regression results with u_(t)_ of the constant-c model. ***C,*** Trial-locked time courses of the brain signals of decision uncertainty in the dACC and the insula. The red and black symbols are the fMRI responses averaged within an ROI and the regression coefficients of u_(t)_ that is estimated by the NBMC, respectively. The significant regression coefficients are marked by the filled circles (GLMM, P < 0.05) and the filled triangles (GLMM, P < 10^−5^). Error bars are 95% GLMM CIs of regression coefficients. ***D,*** Results of the multiple linear regressions of the fMRI responses in the dACC and the insula onto the congruencies of current decisions with stimuli and past decisions. The regression coefficients for the brain signals associated with u_(t)_ (black symbols) are juxtaposed with the regression coefficients for the model simulation of u_(t)_ (green symbols). The significant regression coefficients (GLMM, P < 0.05) for the brain signals are marked by the hexagons, the 95% CIs of which did not include zero. For clarity, only the averages of 95% CIs for the mean of the regression coefficients for the brain signals are shown as the vertical black error bars.

Importantly, when we carried out the regression analysis on those fMRI responses as we did on the RT data, the dACC responses were significantly regressed onto the congruence of d_(t)_ past stimuli but not onto the congruence of d_(t)_ with past decisions (Fig. 6*D*), which corresponds with the history effects of past stimuli on u_(t)_ that are implied by the NBMC. However, the history effect was not statistically significant in the Ins, which indicates that the dACC’s activity most faithfully reflects the changes in decision uncertainty as predicted by the NBMC.

Furthermore, when the values of u_(t)_ that were estimated by the constant-c model were used instead of those estimated by the NBMC, we could not identify any cortical activities that are significantly correlated with u_(t)_, including the dACC and Ins (Fig. 6*B*). This suggests the critical usefulness of the c-inference algorithm in searching for the decision-uncertainty signal in the brain activity of humans engaged in a classification task.

### Dissecting stimulus and criterion contributions to the trial-to-trial variability of classification (P3)

According to the s-inference (Equation 1) and c-inference (Equation 2) algorithms, s_(t)_ and c_(t)_ are inferred based on independent evidences (current and past sensory measurements, respectively) and thus vary independently across trials. According to the deduction algorithms (Fig. 3*E*), the variabilities of d_(t)_ that originate from c_(t)_ and from s_(t)_ can be dissected by regressing the variability of d_(t)_ under a fixed value of s_(t)_ onto the variability of c_(t)_ (left panel of Fig. 7*A*) and by regressing the variability of d_(t)_ under a fixed value of c_(t)_ onto the variability of s_(t)_ (right panel of Fig. 7*A*). This implication of the NBMC captures the third property of classification (P3). Rephrasing in terms of the apple-classification scenario, the NBMC specifies how much of the variability of classification is ascribed to the farm from which an apple came (Fig. 2*B*) and to the farm from which a farmer came (Fig. 2*C*).

**Figure 7.**
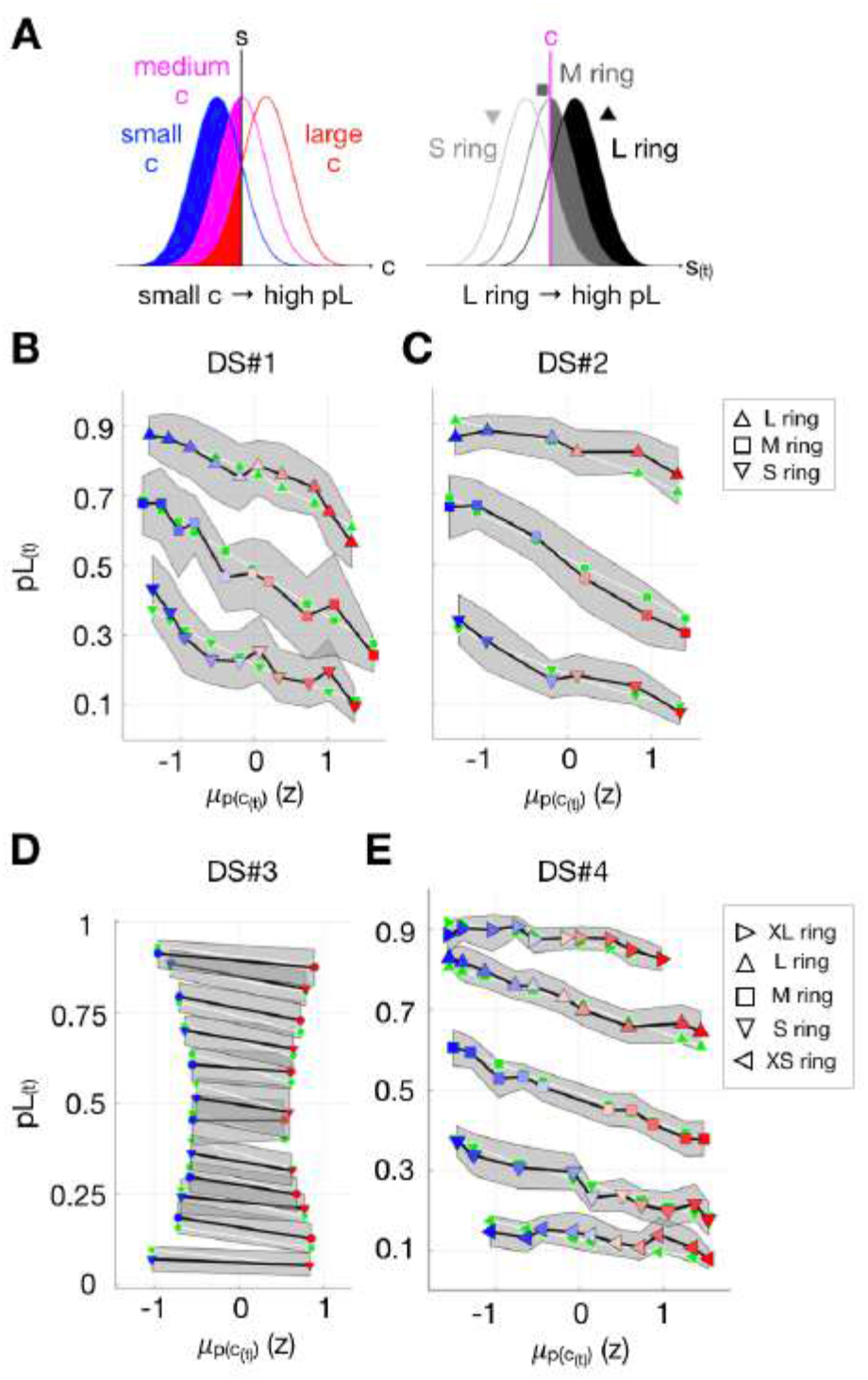
Dissecting stimulus and classification criterion contributions to the variability of decision fraction. ***A,*** A schematic illustration of the respective contributions of c and s to the variability of pL. Left, c varies while s is fixed. Right, s varies while c is fixed. The shaded areas represent the fractions of s being larger than c (i.e., pL). ***B-E,*** Comparisons of pL between the Bayesian and human observers. The average pLs of the Bayesian observers (green symbols) and the humans (blue to red symbols) are plotted against the binned model estimates of c. The data falling on each of the continuous lines are the pLs from the same ring size. Trials were binned by the mean of the inferred criteria 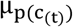, as indicated by the symbols’ color that varies in a scheme similar to that in the left panel of *A*. The shaded gray areas demarcate the 95% bootstrap confidence intervals of data means.

To check how well the NBMC’s dissection of s_(t)_ and c_(t)_ contributions to the variability of d_(t)_ captures the corresponding dissection in the human data, we proceeded as follows. For each stimulus condition, we first binned the trials according to the simulated values of c_(t)_ and calculated the fractions of ‘large’ decision (pL) of the Bayesian observers at those bins (green symbols in Fig. 7*B-E*). Then, we calculated the pL of the human observers at each of all the above bins (blue or red symbols in Fig. 7*B-E*). At all of the bins, the pLs of the humans matched those of the Bayesian observers. Qualitatively, we regressed the human decisions onto current stimulus and the criterion inferred by the NBMC, as follows: 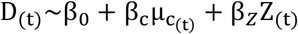. As predicted by the NBMC, the decisions of the humans were concurrently regressed onto c_(t)_ negatively (multiple logistic regressions; β_c_(P) = −0.59 (2.3×10^−20^), −0.54 (2.5×10^−29^), −0.25 (5.4×10^−19^), and −0.35 (3.9×10^−46^) for DS#1~4) and onto the current stimulus positively (β_Z_ P = 1.08 (1.8×10^−35^), 1.36 (9.9×10^−87^), 1.46 (10^−323^), and 1.44 (6.9×10^−71^) for DS#1~4) for all of the data sets. Quantitatively, for all of the data sets, the Bayesian pLs fell within the 95% confidence intervals of the human pLs defined at most of the bins inspected.

Similar to the variability of d_(t)_, the variability of u_(t)_ that originates from c_(t)_ and from s_(t)_ can be dissected by regressing the variability of u_(t)_ under a fixed value of s_(t)_ onto the variability of c_(t)_ (left panel of Fig. 8*A*) and by regressing the variability of u_(t)_ under a fixed value of c_(t)_ onto the variability of s_(t)_ (right panel of Fig. 8*A*). However, unlike d_(t)_, which has a monotonic relationship with v_(t)_, the relationship of u_(t)_ with v_(t)_ is contingent on decisions, being positive for ‘small’ decisions and negative for ‘large’ decisions. For this reason, to check the correspondence in s_(t)_ and c_(t)_ contributions to the variability of u_(t)_ between the Bayesian observers and the humans, we followed the procedure identical to that used for d_(t)_ except that data bins were further conditioned on current decisions. At most of the bins, the RTs and dACC responses of the humans, which can be considered as the behavioral and neural correlates of u_(t)_ as mentioned earlier, matched the u_(t)_ values of the Bayesian observers. Qualitatively, we regressed the RTs in all the data sets, and the dACC responses in DS#1 as well, onto c_(t)_ with stimulus sizes, as follows: 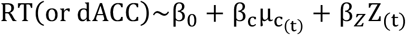. As predicted by the NBMC (Fig. 8*B-F*), β_c_ was positive for ‘large’ decisions (β_c_(P) = 0.12 (2.5×10^−5^), 0.16 (2.5×10^−10^), 0.085 (5.1×10^−8^), and 0.051 (1.7×10^−7^) for RT in DS#1~4 and 0.084 (5.7×10^−4^) for dACC in DS#1) and negative for ‘small’ decisions (β_c_(P) = −0.055 (0.053), −0.21 (2.4×10^−15^), −0.038 (0.013), and −0.058 (9.5×10^−6^) for RT in DS#1~4 and −0.059 (0.020) for dACC in DS#1). Also, β_Z_ was negative for ‘large’ decisions (β_Z_(P) = −0.19 (6.9×10^−15^), −0.21 (2.4×10^−15^), −0.23 (6.0×10^−50^), and −0.097 (2.8×10^−7^) for RT in DS#1~4 and −0.054 0.053 for dACC in DS#1) and positive for ‘small’ decisions (β_Z_(P) = 0.20 (3.5×10^−16^), 0.28 (7.9×10^−28^), 0.21 (3.5×10^−42^), and 0.12 (1.0×10^−12^) for RT in DS#1~4 and 0.11 (5.8×10^−5^) for dACC in DS#1). Quantitatively, for all of the data sets, the Bayesian simulations of u ! fell within the 95% confidence intervals of the human RT and dACC data defined at most of the bins inspected.

**Figure 8.**
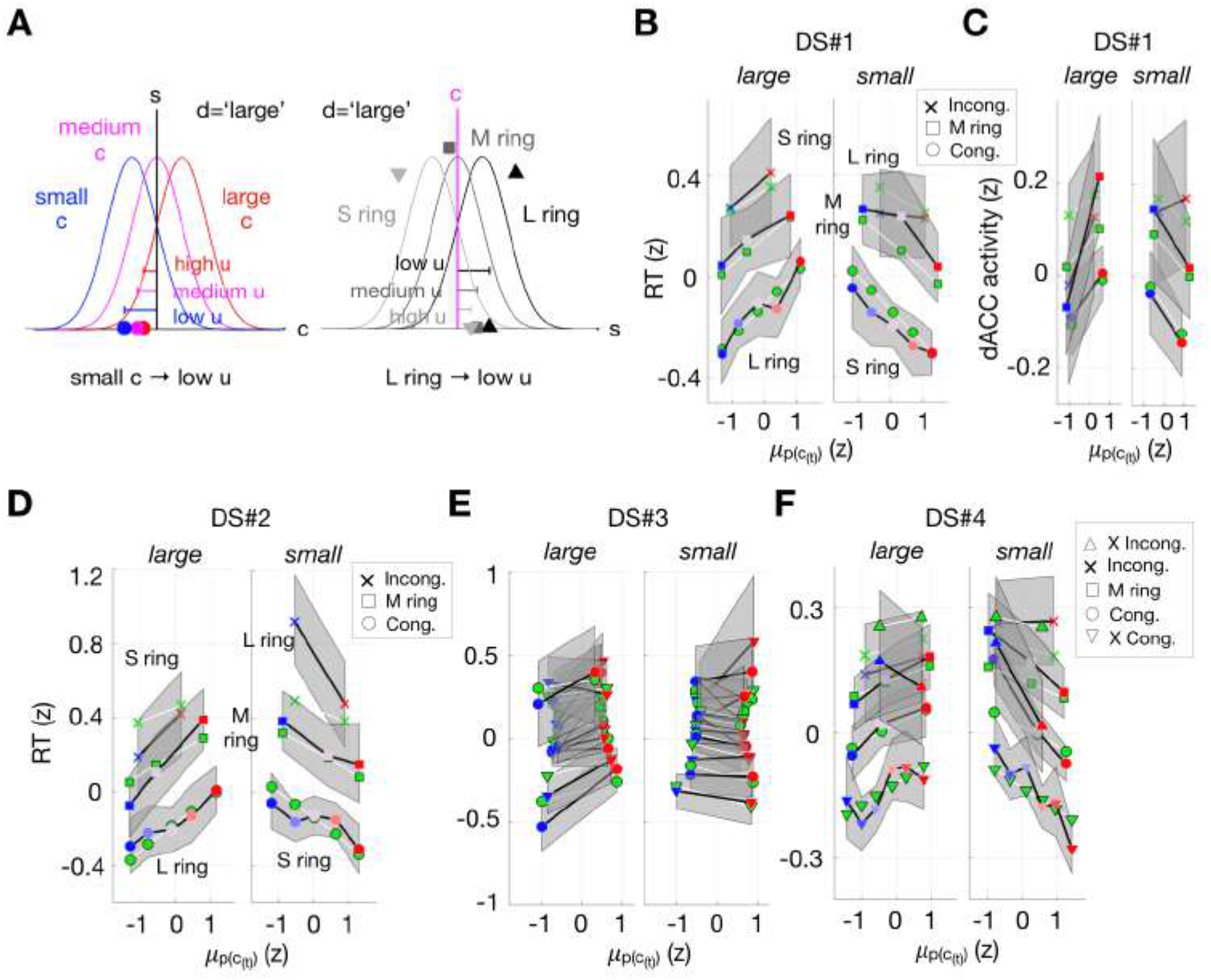
Dissecting stimulus and classification criterion contributions to the variability of decision uncertainty. ***A,*** A schematic illustration of the respective contributions of c and s to the variability of u for ‘large’ decision. Left, c varies while s is fixed. Right, s varies while c is fixed. The symbols on the horizontal axes demarcate the means of c (left panel) and s (right panel) values that yield the ‘large’ decision. For a given condition, the distance between c and s, which is indicated by the length of the horizontal bars, determines the degree of decision uncertainty. ***B-E,*** Comparisons of the decision uncertainty u_(t)_ of the Bayesian observers and the RTs (*B*, *D*-*F*) or the dACC responses (*C*) of the human observers. The average u_(t)_ of the Bayesian observers (green symbols) and the average RTs or dACC responses of the humans (blue to red symbols) are plotted against the binned model estimates of c. The data falling on each of the continuous lines are the data from the same ring size. Trials were binned by the mean of inferred criteria 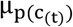, as indicated by the symbols’ color that varies in a scheme similar to that in the left panel of *A*. The shaded gray areas demarcate the 95% bootstrap confidence intervals of data means.

### The relativity of classification (P4)

According to the NBMC, the decision variable v_(t)_, which determines d_(t)_ and u_(t)_, is defined as the difference between s_(t)_ and c_(t)_. This implication realizes the relativity of classification, the last property that the NBMC aims to capture (P4): classification is not governed by the absolute magnitude of stimuli but by the relative magnitude of s to c, as intuited with the different pairings of the farms from which apples and farmers come in the apple-classification scenario. Figure 9A and 10A illustrate how the NBMC captures the relativity of classification in terms of pL_(t)_ and u_(t)_, respectively: pL_(t)_ and u_(t)_ should both be distinguishable as the differences between s_(t)_ and c_(t)_ are unmatched even though physical stimuli are the same (the left two panels of Fig. 9*A* and Fig. 10*A*) and indistinguishable as long as the differences are matched even though physical stimuli are different (the right two panels of Fig. 9*A* and Fig. 10*A*).

**Figure 9.**
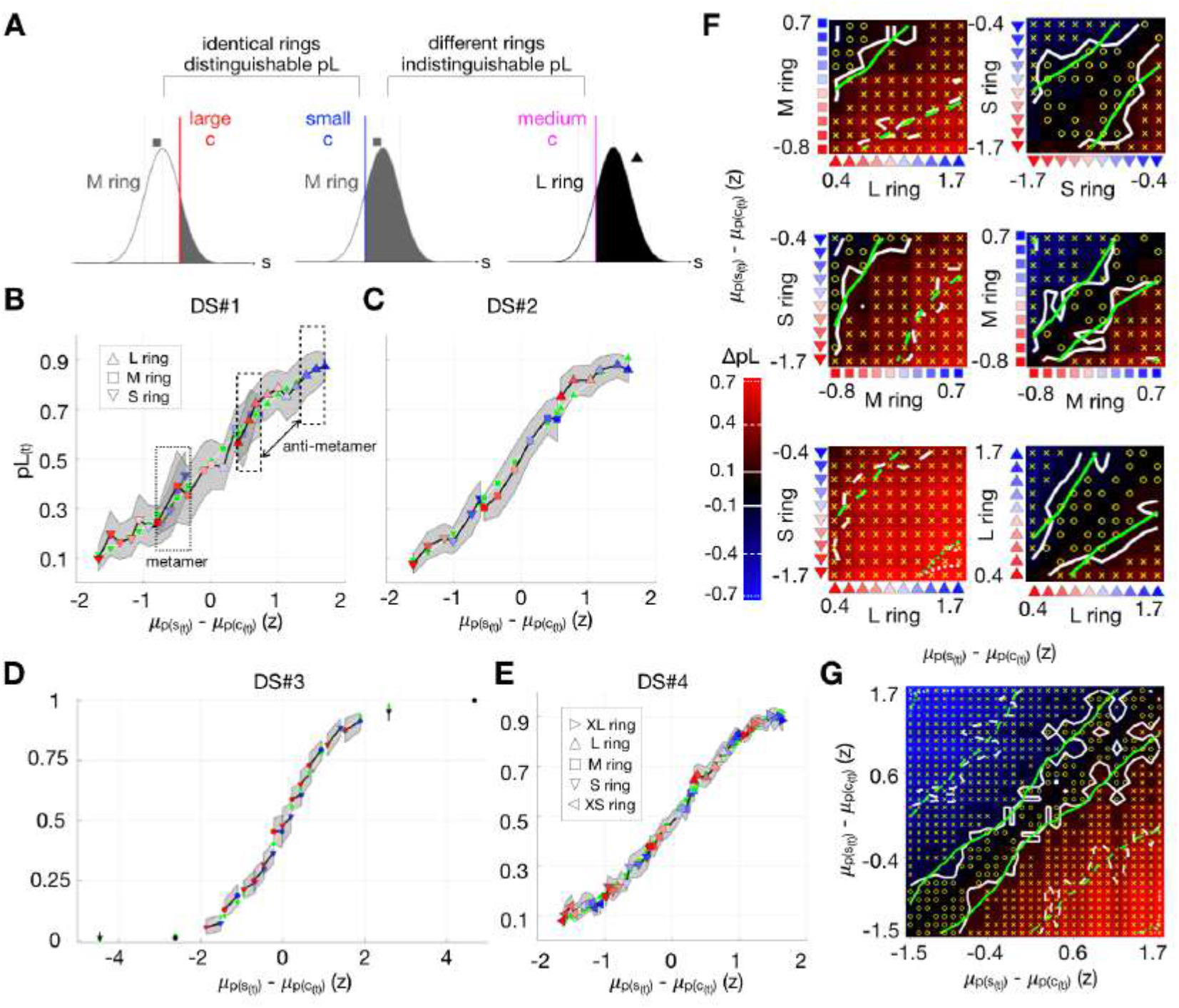
Relativity of classification in decision fraction. ***A,*** A schematic illustration of ‘anti-metameric’ and ‘metameric’ trial pairs in decision fraction. The curves represent the distributions of s for given ring sizes. The light gray vertical lines demarcate the means of s for the S, M, and L-size rings. The vertical lines in color represent the locations of c. The shaded areas represent pLs, where s is greater than c. In the ‘anti-metameric’ pair (left two panels), the pLs differ substantively despite identical stimuli due to a large difference in c. In the ‘metameric’ pair (right two panels), the pLs do not differ despite different stimuli because the locations of c relative to the distribution of s do not differ. ***B-E,*** Continuous and monotonic increases in pL_(t)_ with increasing differences between s and c over different stimulus conditions. The pLs of the Bayesian observers (green symbols) and the humans (non-green symbols) are plotted against the differences between s and c. The color spectrum from blue to red represents the increasing size of c within each stimulus condition. The symbol shapes represent different stimulus conditions (but only two different shapes were used alternatingly to distinguish many different ring sizes for DS#3). Shaded areas, 95% bootstrap confidence intervals of the means of data. ***F-G,*** Statistical demonstration of the relativity in DS#1 by using Bayes factor. ***F,*** For all of the possible pairs of the trial bins in DS#1 (shown in *B*), the differences in pL_(t)_ were calculated and represented by color gradients within 6 separate grid panels. For each grid panel, the columns and rows of trial bins are marked by the same symbols and colors used in *B*. ‘x’ and ‘o’ markers indicate pL_(t)_ of the bin pairs that are significantly distinguishable (Bayes factor > 3) and indistinguishable (Bayes factor < 1 3), respectively. The dotted, dashed and solid lines are the iso-ΔpL lines with ΔpL = ±0.7, ±0.4, ±0.1, respectively. The green and white lines are the iso-ΔpL lines defined by the Bayesian simulations and the human data, respectively. ***G,*** A unified grid of ΔpL merged across stimuli. The format is identical to that used in *F*.

**Figure 10.**
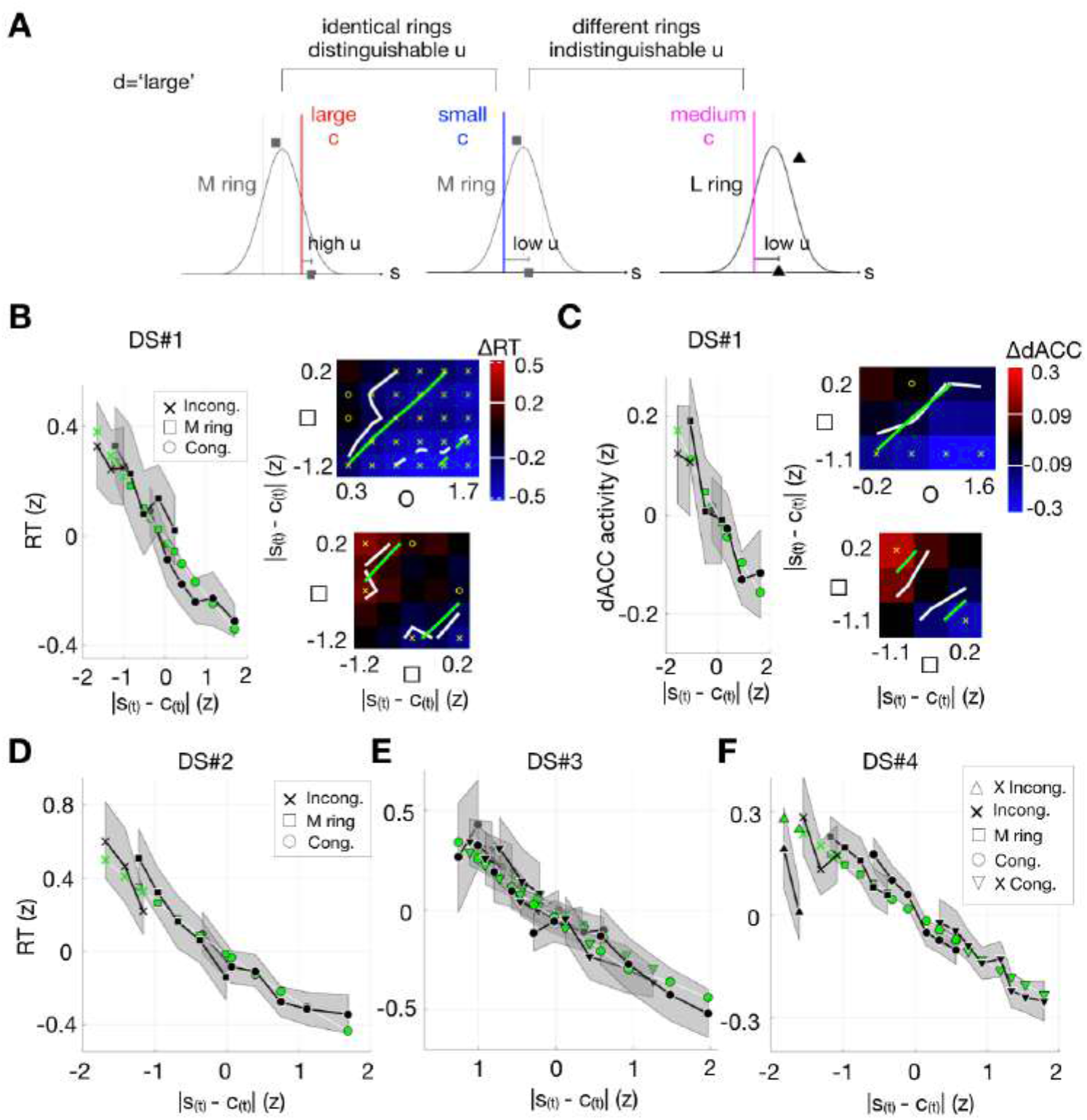
Relativity of classification in decision uncertainty. ***A,*** A schematic illustration of ‘anti-metameric’ and ‘metameric’ trial pairs in decision uncertainty. The curves represent the distributions of s for given ring sizes. The light gray vertical lines demarcate the means of s for the S, M, and L-size rings. The vertical lines in color represent the locations of c. The length of the horizontal bars represents the mean degree of decision uncertainty (u_(t)_), when a current decision is ‘large’. In the ‘anti-metameric’ pair (left two panels), the u_(t)_s differ substantively despite identical stimuli due to a large difference in c. In the ‘metameric’ pair (right two panels), the u_(t)_s do not differ despite different stimuli because the locations of c relative to the distribution of s do not differ. ***B-F,*** Continuous and monotonic increases in u_(t)_, RT, or dACC activity with increasing differences between s and c over different stimulus conditions. The u_(t)_ of the Bayesian observers (green symbols) and the RTs or dACC responses of the humans (black symbols) are plotted against the differences between s and c. The symbol shapes represent different stimulus conditions (but only two different shapes were used alternatingly to distinguish many different ring sizes for DS#3). Shaded areas, 95% bootstrap confidence intervals of the means of data. ***B-C,*** The right panels show the results of the Bayes factor analysis on the RT (*B*) and dACC (*C*) data for DS#1. For the possible pairs of the trial bins in Cong.-M ring condition (top) or M ring-M ring condition (bottom), the differences in RT (*B*) or in dACC activity (*C*) were calculated and represented by color gradients. For each grid panel, the columns and rows of trial bins are marked by the same symbols and colors used in the left panels. ‘x’ and ‘o’ markers indicate RTs or dACC responses of the bin pairs that are significantly distinguishable (Bayes factor > 3) and indistinguishable (Bayes factor < 1 3), respectively. The dashed and solid lines in *B* are the iso-ΔRT lines with ΔRT= ±0.5, ±0.2, respectively. The solid lines in *C* are the iso-ΔdACC lines with ΔdACC = ±0.09. The green and white lines are the iso-ΔRT or iso-ΔdACC lines defined by the Bayesian simulations and the human data, respectively.

To evaluate how well the NBMC captures the relativity of human classification in pL_(t)_, we checked the correspondence between the Bayesian observers and the humans in the following two model implications. First, because what governs classification is ‘s relative to c’, pL_(t)_ should vary only as a single, monotonic function of the difference between s_(t)_ and c_(t)_. When trials were sorted by the differences between the simulated values of s_(t)_ and cA, the pL_(t)_ values of the humans constituted single, seamless psychometric curves for all of the data sets (red or blue symbols in Fig. 9*B-E*), and most of the pL_(t)_ values of the Bayesian observers (green symbols of Fig. 9*B*-E) fell within the 95% confidence intervals of the human pL_(t)_.

Second, again because what governs classification is ‘s relative to c’, pL_(t)_ should become ‘metameric’ between the trials in which a physical difference between stimuli is compensated by a counteracting difference in c_(t)_ (e.g., dotted box in Fig. 9*B*) or ‘anti-metameric’ between the trials in which a physical sameness between stimuli is accompanied by a substantial difference in c_(t)_ (e.g., dashed boxes in Fig. 9*B*). These predictions were statistically confirmed by the Bayes factor analysis across all the possible pairs of trial bins (see Methods for the procedure for the Bayes factor analysis). The distributions of the “significantly indistinguishable” trials (‘o’ markers in the left columns of Fig. 9*F*; Bayes factor < 1/3) and the “significantly distinguishable” trials (‘x’ markers in the right columns of Fig. 9*F*; Bayes factor > 3) coincided with the metameric relations predicted by the NBMC. Figure 9G, which puts together the panels in Figure 9F, demonstrates that the NBMC readily captures the relativity of classification in a single currency, the difference between s_(t)_ and c_(t)_.

The RT and dACC data, though being noisier than the decision fraction data, also supported the Bayesian-human correspondences in u_(t)_ regarding the relativity of classification. When trials were sorted by the differences between the simulated values of s_(t)_ and c_(t)_, the human RT (black symbols in Fig. 10*B, D-F*) and dACC (black symbols in Fig. 10*C*) responses constituted single, seamless psychometric curves for all of the data sets, and most of the u_(t)_ values of the Bayesian observers (green symbols of Fig. 10*B*-F) fell within the 95% confidence intervals of the human RT (black symbols in Fig. 10*B-E*). Furthermore, the pairs of trials in which the human RT and dACC responses became significantly metameric (‘o’ markers in the upper right panels of Fig. 10*B* and Fig. 10*C*; Bayes factor < 1/3) and significantly anti-metameric (‘x’ markers in the bottom right panels of Fig. 10*B* and Fig. 10*C*; Bayes factor > 3) were found.

### Brain signals of inferred stimulus, criterion, and decision variable

The Bayesian observer of the NBMC forms the five variables that are causally related with one another, s_(t)_, c_(t)_, d_(t)_, and u_(t)_. So far, to verify the implications of the NBMC, we have shown that its simulation of d_(t)_ and u_(t)_ qualitatively and quantitatively captures the core properties of classification (P1~4) that are expressed in the pL and RT (or dACC activity) measurements acquired from the human observers. As indicated by the causal structure of the variables (Fig. 3*E*), the simulated values of d_(t)_ and u_(t)_ were derived from their parent variables, s_(t)_, c_(t)_, and v_(t)_, the trial-to-trial latent states of which were predicted by the NBMC. This allows us to examine the fMRI responses of the human observers in search of the brain activities that represent s_(t)_, c_(t)_, and v_(t)_. Such brain activities should satisfy the following two requirements: first (‘correlation’ requirement), for each brain signal, its trial-to-trial states must be correlated with those of its target variable; second (‘causality’ requirement), its correlations with the other remaining (non-target) variables in the NBMC must be consistent with the causal relationships of its target variable with the remaining variables.

Initially, we identified the candidate brain regions within which activity patterns satisfy the first ‘correlation’ requirement using the multivoxel pattern analysis (MVPA) in conjunction with a searchlight technique, which is known to be highly sensitive to detect brain signals in local patterns of population fMRI responses (Kriegeskorte et al., 2006). However, this approach is limited in specificity: successful decoding of a variable from certain patterns of brain activity does not necessarily indicate the representation of that variable per se because such decoding is also possible even when those patterns of activity indeed represent any other variables that are correlated with the variable of interest.

To address this specificity problem, we imposed the second ‘causality’ requirement on the candidate brain regions. Specifically, we deduced an exhaustive set of regressions (Fig. 11*D-F*) that are implied from the causal relationships of a target variable with other variables (Fig. 11*A-C*), and tested whether a candidate brain signal is consistent with the entire set of regressions. For example, to claim that a candidate brain signal of c satisfies the ‘causality’ relationship between c and v, we must show that that the c decoded from the brain signal is (i) negatively regressed onto v (the 5th regression in Fig. 11*D*) but (ii) not negatively regressed onto v any more when the causal influence of c is removed in v (the 6th regression in Fig. 11*D*), which are implied by the negative causality between c and v (the top left arrow in Fig. 11*A*; the histogram of correlation coefficients between c and v in Fig. 11*B*). Additional regressions can be further deduced from the causal relationship of c with the remaining variables (s, u, d, *Z*), resulting in a total of 13 regressions for c (Fig. 11*D*). Likewise, we deduced 13 and 16 regressions for s and v, respectively (Fig. 11*D-F*; see Methods for detailed definitions of the regressions).

**Figure 11.**
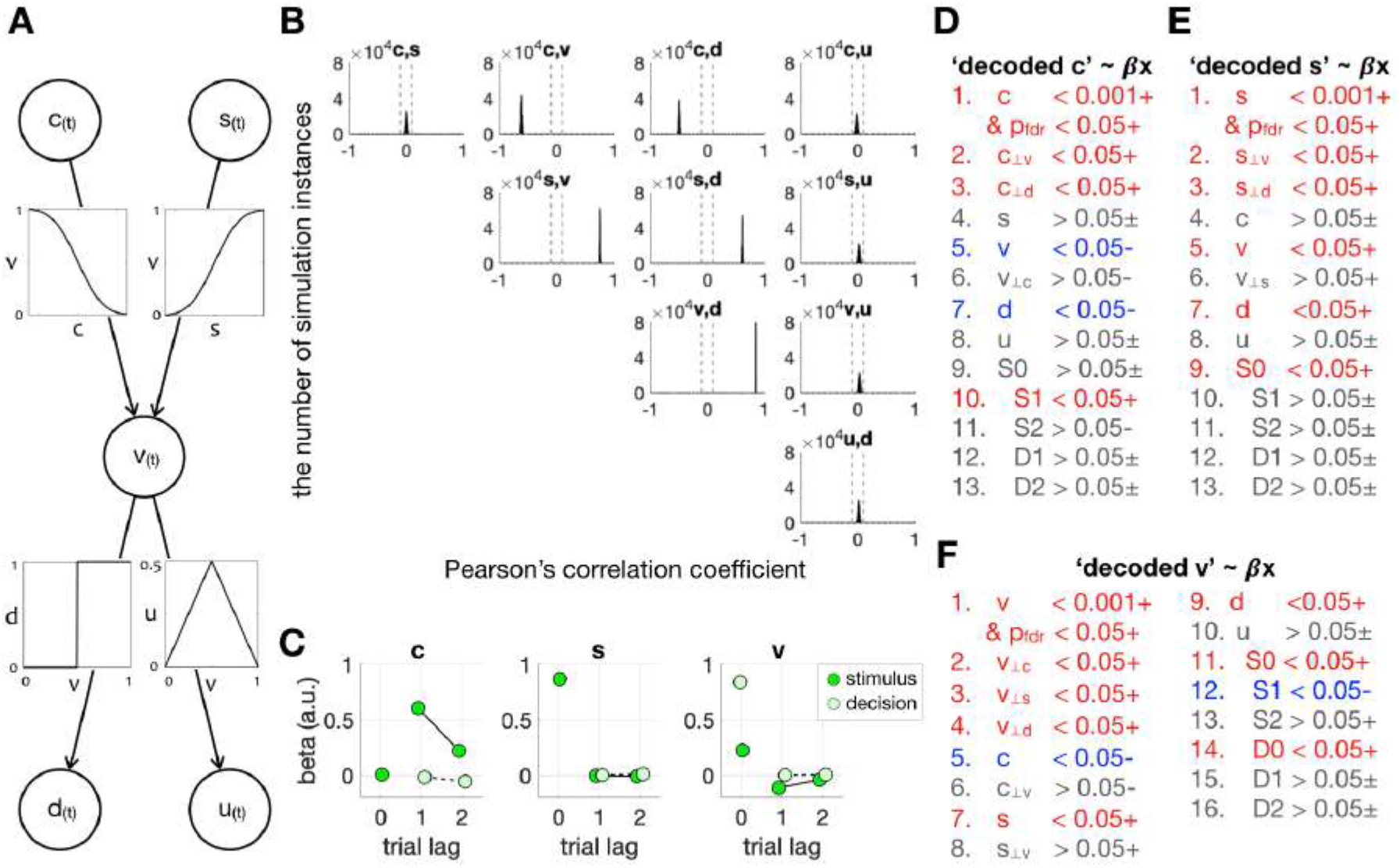
Deduction of regressions for c_(t)_, s_(t)_, and v_(t)_ from the causal structure of variables in the NBMC. ***A,*** A graphic illustration of the causal structure of variables. Each arrow and an accompanying inset plot represent the direction and function of the causality between two variables, respectively. ***B,*** Histograms of simulated correlation coefficients between the variables. For each possible pair of the variables, the distribution of Pearson’s correlation coefficients across many repeated simulations (see Methods for the simulation procedure) was plotted as a histogram. The gray dashed lines demarcate ±0.1 Pearson’s correlation coefficients. ***C,*** Multiple linear regressions of the simulated values of c_(t)_, s_(t)_, and v_(t)_ onto the stimuli and decisions on current and past trials. ***D-F,*** The lists of regressions that should be satisfied by c_(t)_, s_(t)_, and v_(t)_. The regressions onto the other latent variables (1~8th, 1~8th, 1~10th regressions for c_(t)_, s_(t)_, and v_(t)_, respectively) were derived from the simulated correlation analyses depicted in *B* while the regressions onto stimuli and decisions (9-13th, 9~13th, 11~16th regressions for c_(t)_, s_(t)_, and v_(t)_, respectively) were derived from the simulated multiple regression analyses depicted in *C*. The meanings of symbols, numbers, and colors used for describing the regressions are as follows: ‘+’, ‘−’, and ‘±’, indicate that the significance test is one-tailed right, one-tailed left, and two-tailed, respectively; numbers with decimal points, threshold p-values for GLMM regression (p_fdr_, FDR-corrected p-values); ‘<’ and ‘>’, significant and non-significant regressions, respectively; ‘A_⊥B_’, residuals of A from the linear regression onto B; ‘D#’ and ‘S#’, decision and stimulus on a #-back trial; red, blue, and gray are positive, negative, and non-significant regressions, respectively.

Six brain regions satisfied both of the two requirements (Table 3; Fig. 12*A*). The signal of c appeared in three different regions at different time points relative to stimulus onset, as follows: a region in the inferior parietal lobe at 1.1 s (IPL_c1_), followed by two regions in the superior temporal gyrus at 3.3 and 5.5 s (STG_c3_, STG_c5_). The signal of s appeared both in the dorsolateral prefrontal cortex at 3.3 s (DLPFC_s3_) and in the cerebellum at 5.5 s (Cereb_s5_). The signal of v appeared in the left superior temporal gyrus at 5.5 s (STG_v5_).

**Table 3.** Specifications of the regions in which the brain signals of c_(t)_, s_(t)_, and v_(t)_ were identified.

| Cortical area | Decoded variable | Informative time from stimulus onset (s) | Abbreviated on of the decoded variable | Contiguous searchlight centers (N) | Peak searchlight |  |
| --- | --- | --- | --- | --- | --- | --- |
|  |  |  |  |  | MNI coordinate | GLMM p-value of |
| Left inferior parietal lobe | $c_{(t)}$ | 1.1 | $IPL_{c1}$ | 15 | [-54, -27, 48] | $8.8 \times 10^{-8}$ |
| Left superior temporal gyrus | $c_{(t)}$ | 3.3 | $STG_{c3}$ | 13 | [-45, -30, 9] | $5.7 \times 10^{-7}$ |
| Left superior temporal gyrus | $c_{(t)}$ | 5.5 | $STG_{c5}$ | 18 | [-66, -21, 9] | $2.3 \times 10^{-7}$ |
| Left dorsolateral prefrontal cortex | $s_{(t)}$ | 3.3 | $DLPFC_{s3}$ | 33 | [-51, 27, 24] | $1.9 \times 10^{-7}$ |
| Right cerebellum | $s_{(t)}$ | 5.5 | $Cereb_{s5}$ | 36 | [36, -63, -21] | $1.4 \times 10^{-6}$ |
| Left superior temporal gyrus | $v_{(t)}$ | 5.5 | $STG_{v5}$ | 15 | [-60, -9, 12] | $4.9 \times 10^{-8}$ |

**Figure 12.**
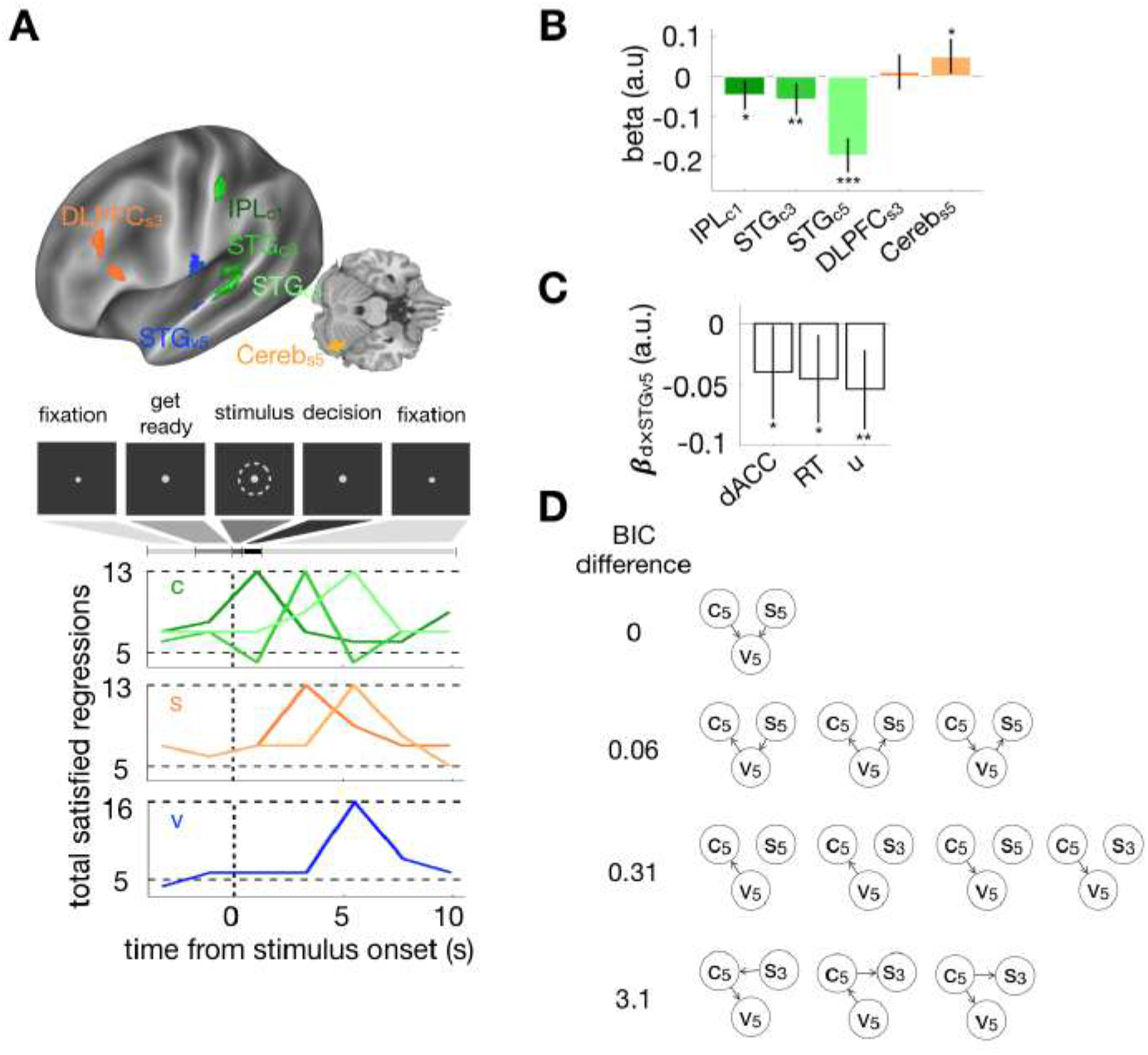
The brain signals of c_(t)_, s_(t)_, and v_(t)_. ***A,*** Top, the six loci of the brain signals that satisfied all the regressions required by the causal structure of variables in the NBMC are overlaid on the template brain. See Table 3 for the detailed specifications for these loci. Bottom, the numbers of the satisfied regressions are plotted as a function of time relative to stimulus onset for the c, s, and v signals. ***B***, Coefficients of multiple regressions of the brain signal of v onto the brain signals of c and s. ***C***, Coefficients of the ‘decision × brain signal of v’ term in the regression models, in each of which either the brain signal of u (dACC_u5_), the RT data, or the model estimates of u, was regressed onto the brain signal of v (STG_v5_) and decisions. ***B***-***C***, *, P < 0.05; **, P < 0.01; ***, P < 0.001. Error bars, 95% GLMM confidence intervals of the mean coefficients. ***D***, The causal networks sorted by their BIC scores relative to the network that best explains the concurrent variabilities between the brain signals of c_(t)_, s_(t)_, and v_(t)_.

We also confirmed that the concurrent trial-to-trial variabilities of the decoded variables from the brain signals were consistent with the causal structure defined by the NBMC, as follows. First, the interplays between the decoded c, s, and v were well captured by a multiple regression model, STG_v5_~β_s3_DLPFC_s3_ + β_s5_Cereb_c_s5 + β_c1_IPL_c1_ + β_c3_STG_c3_ + β_c5_STG_c5_, which was derived from the ‘common-effect’ structure of causal relationships between the variables (c → v ← s; the top edges of Fig. 11*A*). The regression coefficients matched the causal functions in sign for all of the five brain signals and were all significant except for DLPFC_s3_ (β_s3_ = 0.011, P = 0.55; β_s5_ = 0.047, P = 0.013; β_c1_ = −0.046, P = 0.0088; β_c3_ = −0.056, P = 0.0037; β_c5_ = −0.15, P = 1.9×10^−17^; Fig. 12*B*). Second, the decision-dependent interplay between v and u signals was successfully captured by an interactive regression model, dACC_u5_~β_v_STG_v5_ + β_d_d + β_d×v_d×STG_v5_, which was derived from the ‘common-cause’ structure of causal relationships between the variables (d ← v → u; the bottom edges of Fig. 11*A*). Critically, because the sign of the causal function from v to u depends on the value of d, the NBMC predicts the interaction between d and STG_v5_. As predicted, dACC_u5_ was significantly regressed onto d×STG_v5_ (β_d×v_ = −0.040, P = 0.04; the left bar in Fig. 12*C*). The causal structure of ‘d ← v → u’ was further supported when the RT data (β_d×v_ = −0.045, P = 0.013) or the model estimates of decision uncertainty u_(t)_(β_d×v_ = −0.054, P = 0.0012), instead of dACC_u5_, were regressed onto d and STG_v5_ (the middle and right bars in Fig. 12*C*).

Finally, as a data-driven approach that complements the model-driven regression analyses performed above, we calculated the Bayesian Information Criterion (BIC) for each of the possible causal structures between the brain signals of c, s, and v (Scutari, 2009); see Methods for BIC calculation). The outcomes of BIC evaluation were consistent with the NBMC in the following two aspects. First, out of the 162 possible causal graph structures, the smallest (best) BIC value was found for ‘c → v ← s’ (Fig. 12*D*). This causal graph of ‘common effect’ is exactly the one prescribed by the NBMC (Fig. 11*A*). Second, note that one critical outcome that falsifies the NBMC is the ‘significant’ existence of any graph that includes causal arrows between c and s because the NBMC is built upon the assumption that c and s are the random variables that are independent from one another. To evaluate the statistical significance of those graphs, we followed the statistical convention that any pairs of hypotheses with the BIC differences greater than 2 are considered to be sufficient to conclude significant differences (Kass and Raftery, 1995). With this critical value of statistical significance (BIC>2), we found that none of the graphs with the causal arrows between c and s could pass the criterion. The closest group of the graphs (shown at the bottom of Fig. 12*D*) was well apart from the winner graph, by more than the BIC value of 3.

In sum, the outcomes of the fMRI data analyses indicate that there exist the brain activities that respectively represent the key latent variables of the NBMC (c, s, and v) and concurrently vary in the ways consistent with the causal relationships of those variables with themselves and the other variables in the NBMC.

### Different history effects in classification and categorization

Although “classification” and “categorization” have been used interchangeably, as footnoted in Introduction, the two tasks should not be confused with each other (Jacob, 2004) since they fundamentally differ in statistical structure. Whereas classification requires judging which side of a single distribution (i.e., ‘which class’) a given entity belongs to relative to a criterion (Fig. 13*A*), categorization requires judging which of two distributions (i.e., ‘which category’) a given entity belongs to (Fig. 13*B*). In the context of the apple-sorting scenario, classification is to decide whether a given apple is smaller or larger than the typical size of apples while categorization is to decide whether a given apple is from the B farm or from the F farm.

**Figure 13.**
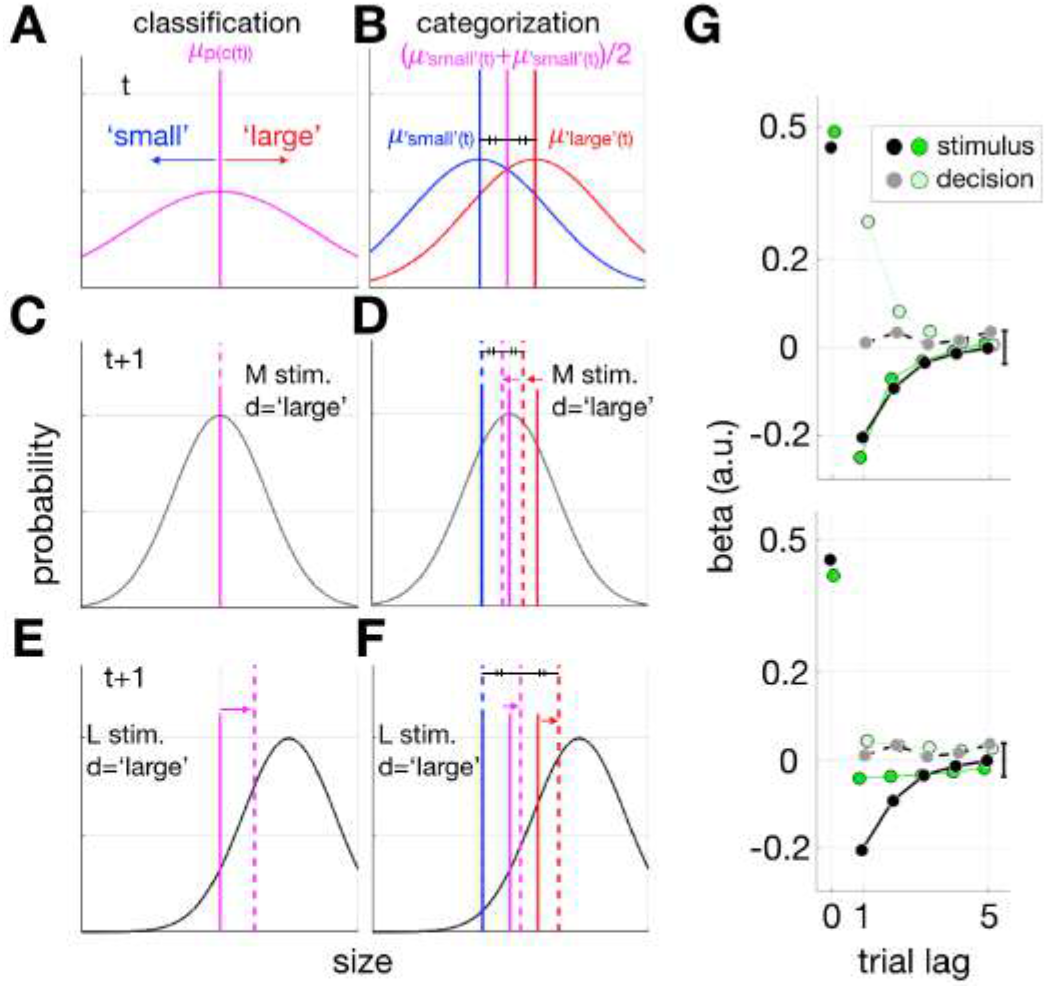
History effects predicted by the categorization model. ***A-F,*** Schematic illustrations of how the criterion and decision boundary are updated in the classification (*A*, *C*, *E*) and categorization (*B*, *D*, *F*) models, respectively. ***A-B,*** Distributions of stimuli assumed in the classification (*A*) and categorization (*B*) models. In the classification model, a stimulus is classified into ‘small’ or ‘large’ according to their size relative to the criterion (dashed line) within a single distribution of stimuli (purple curve). In the categorization model, a stimulus is categorized into ‘small’ or ‘large’ according to which ‘category’ distributions of stimuli (blue and red curves for ‘small’ and ‘large’ categories, respectively) it belongs to. Thus, when the two distributions are equal in shape, the effective decision boundary (dashed purple line) is prescribed as the arithmetic mean of the means of the two category distributions (dashed blue and red lines). ***C-D,*** Effects of past decisions on classification criterion (*C*) and categorization boundary (*D*). Gray curves represent distributions of noisy sensory measurements associated with a medium-size stimulus. When a ‘large’ decision was made on a medium-size stimulus in a past trial, the classification criterion (solid purple line in *C*) remains the same whereas the categorization boundary (solid purple line in *D*) is shifted toward the small side as the ‘large’ category distribution (solid red line in *D*) is shifted toward the small side. ***E-F,*** Effects of past stimuli on classification criterion (*E*) and categorization boundary (*F*). Gray curves represent distributions of noisy sensory measurements associated with a large-size stimulus. When a large-size stimulus was presented in a past trial, the classification criterion (solid purple line in *E*) and the categorization boundary (solid purple line in *F*) are both shifted toward the small side. ***G,*** Regressions of the simulated decisions of an observer with the categorization-model onto stimuli and past decisions for DS#1. The regression coefficients for the simulated data (green symbols) are juxtaposed with those for the human data (black and gray symbols). The top and bottom panels show the results associated with the two different set of model parameters, the former being best fit to stimulus history effects and the latter to decision history effects.

Due to this difference in statistical structure, classification and categorization predict different history effects. The history effects in classification occur as an observer updates the single, entire distribution of stimuli in her generative model through recent experience of stimuli, and thus the classification criterion (the median of the distribution). Critically, past decisions are not involved at all in the history effects in classification because past decisions do not provide any information about the distribution of stimuli (Fig. 13*C*)—only past experience of stimuli can attractively shift the distribution of stimuli and thus the criterion (Fig. 13*E*). As results, only past stimuli, but not past decisions, bring about negative history effects on current decisions as observed in the current study (Fig. 4).

By contrast, the history effects in categorization occur as an observer selectively updates the two category likelihoods over stimuli in his generative model. Importantly, unlike in classification, ‘past decisions’ contribute to the history effects in categorization by determining which of the two category likelihoods is updated because assigning a stimulus to a certain decision means that the stimulus belongs to the category corresponding to that decision (Fig. 13*D*). On the other hand, like in classification, ‘past stimuli’ attractively shift only that selected category likelihood toward themselves (Fig. 13*F*). As results, in categorization, ‘past decisions’ tend to be repeated on current trials while ‘past stimuli’ tend to repel current decisions. Alternatively, the history effects in categorization can also occur as an observer updates the prior probabilities of the categories in her generative model, as suggested recently (Glaze et al., 2015). Even in this alternative updating, past decisions still contribute to the history effects by determining which of the two category priors is increased (or decreased, depending on other assumptions in the generative model such as ‘hazard’ (Glaze et al., 2015)). In sum, due to the difference in statistical structure, classification and categorization critically differ in history effect in that ‘past decision’ is a critical factor in categorization but not in classification.

To concretely demonstrate the history effects in categorization, we adapted one of recent models of categorization (Norton et al., 2017) to our experiment (see Methods for model specifications) and examined the history effects through model simulations. This particular model, which updates the likelihoods of categories over stimuli and was called the “exponentially weighted moving average” model in Norton et al. (2017), was chosen because it best captured human categorization while exhibiting the history effect of past stimuli similar to that in the current study. When we carried out a regression analysis on the simulated data, as was done to probe the history effects on decisions for the NBMC (Fig. 4), the regression coefficients of past stimuli and past decisions were opposite in sign (negative for past stimuli and positive for past decision) and similar in magnitude (green symbols in Fig. 13*G*). These history effects differed from those observed for the humans in our experiments (black and gray symbols in Fig. 13*G*). This difference could not be resolved by changing the model parameters: the parameter set that best captures the stimulus effect fails to capture the decision effect (top panel of Fig. 13*G*), and vice versa (bottom panel of Fig. 13*G*).

## Discussion

By specifying the statistical structure of classification, we developed the NBMC, in which a normative observer infers the stimulus and criterion from current and past sensory evidences, respectively, and commits to a decision with a varying degree of uncertainty. The NBMC well captured human classification behavior and neural activity by precisely defining the latent process of criterion inference. Furthermore, we demonstrated how the difference in statistical structure between classification and categorization leads to the difference in history effect.

### Normative inference of classification criterion from sensory evidences with memory decay

By characterizing our model as “normative”, we mean that, under the assumption that sensory traces of past stimuli become noisier with memory decay, the optimal way of inferring the center of past stimuli is to probabilistically integrate the likelihoods by taking into account their increasing uncertainties associated with memory decay since such integration minimizes the squared error of inferred estimates (Jaynes, 2003; Körding and Wolpert, 2004). To be sure, for an agent without memory decay, this algorithm is suboptimal because the inferred criterion keeps digressing from the true, fixed criterion by being swayed with recent stimuli. In this sense, the c-inference algorithm proposed here can be considered as an instantiation of the bounded rationality of an agent with limited cognitive resources (Simon, 1955).

### Classification versus categorization

Classification requires learning about a criterion, which is arbitrarily set and divides a distribution of entities into mutually exclusive classes. For example, classifying the roasting degrees of coffee beans into ‘dark-medium’ or ‘light-medium’ requires setting an arbitrary criterion that divides a given distribution of roasting degrees into two exclusive parts. Here, ‘dark-medium’ or ‘light-medium’ is a relative adjective that is assigned to a bean via comparison against an arbitrary criterion.

On the other hand, categorization requires a knowledge about how feature values are distributed for given categories (Glaze et al., 2015; Gold and Shadlen, 2007; Green and Swets, 1966; Maddox, 2002; Norton et al., 2017; Qamar et al., 2013). For example, categorizing the origin of coffee beans into ‘Ethiopia’ or ‘Brazil’ based on color requires knowing about the respective color distributions of the Ethiopian and Brazilian beans. Here, the categorical membership of a given bean is determined by the distributions of color feature intrinsic to the geographical origin of the bean, not by any arbitrary criterion value of color. Therefore, unlike in classification, the color distributions may be overlapped between ‘Ethiopian’ and ‘Brazilian’ beans.

Due to this difference in statistical structure, classification and categorization entail different generative models and inferential processes. As prescribed by the NBMC, the ideal classification is to infer both the criterion (c) that divides the ‘dark-medium’ and ‘light-medium’ classes and the amount of roasting (s) by propagating the information gleaned from past (r) and current sensory (m) measurements of roasting, respectively, and then to commit to a decision by computing the posterior probability that the amount of roasting is greater than the criterion, p(s > c|m, r). By contrast, the ideal categorization is to infer the posterior probabilities of ‘Ethiopia’ and ‘Brazil’ given sensory measurements as p(Ethoipia|m) and p(Brazil|m) and then to judge which category has the higher posterior.

Although the size classification task is performed ideally by the NBMC, human observers could have performed the task by adopting the categorization model because doing so leads to a “satisfactory” level of performance though not the ideal level. However, this seems unlikely because the observers in our study did not exhibit history effects of past decisions on current decisions whereas an observer who adopts the generative model of categorization would exhibit substantive history effects of past decisions irrespective of whether the observer updates the likelihoods or priors of categories. Our results suggest that researchers who develop or use any binary decision-making tasks need to specify whether their task entails classification or categorization and to apply an appropriate model accordingly. We stress that the NBMC should be considered as a computational model for binary classification tasks in which a criterion is not explicitly provided.

### History effects in classification via dynamic criterion update

The history effects in sequential perceptual decision-making have recently received considerable attention. The NBMC offers a normative account for the repulsive effect of past stimuli on classification of a current stimulus by specifying how the Bayesian inference of the criterion is dynamically updated from past sensory evidences. This account is distinct from those offered by previous studies in the following aspects.

First, in categorization tasks, history effects occur as the likelihoods (Norton et al., 2017) or priors (Glaze et al., 2015; Glaze et al., 2018) of categories are updated. As shown in the last section of Results, these categorization-based accounts are different from our classification-based account in that they predict the systematic influences of past decisions on current decisions.

Second, unlike in many previous accounts (Burr and Cicchini, 2014; Dyjas et al., 2012; Fritsche et al., 2017; Glaze et al., 2015; Raviv et al., 2012; Treisman and Williams, 1984), any specific assumption regarding environmental volatility is not required in our account for history effects. Instead, our account is grounded in the assumption that the uncertainty of a past sensory event increases as trials elapse (Gorgoraptis et al., 2011; Zokaei et al., 2015).

Third, as proposed previously (Fritsche et al., 2017; Schwartz et al., 2007; Stocker and Simoncelli, 2006), sensory adaptation might be considered as an alternative origin for the repulsive history effect of past stimuli because an exposure to a specific stimulus would induce a repulsive bias on a population neural representation. However, this account seems unlikely because stimuli were presented very briefly (0.3s) and the history effect of past stimuli sustained more than 26.4s in the current study whereas the neural bias due to sensory adaptation requires a long duration of exposure and lasts for a short period of time (Nakashima and Sugita, 2017; Pavan et al., 2012).

### Brain signals of inferred criterion

The brain signals of c appeared initially at the IPL and then at the STG, in the vicinity of which the signal of v resides. We conjecture that working-memory representations of past stimuli are likely to be formed in the IPL and then are transferred to the STG, wherein they take part in classification of current stimuli. This conjecture is congruous with the literature in several aspects.

Optical inactivation of the posterior parietal cortex (PPC) of rat – a region presumed to be a rat homologue of the primate IPL (Goard et al., 2016; Hanks et al., 2015; Roitman and Shadlen, 2002) – selectively diminished the influence of previous stimuli but left the task performance on current stimuli intact (Akrami et al., 2018), suggesting a critical role of the IPL in encoding previous stimulus information. In addition, perturbing the PPC after current stimulus onset failed to diminish history effects, suggesting that the information of previous stimuli in PPC was likely to be transferred to some brain regions outside the PPC (Hwang et al., 2017). On the other hand, the c signal at the STG seems consistent with its suggested roles in coordinating the spatial reference frame (Karnath, 2001; Karnath et al., 2001; Karnath et al., 2002), in that c in our task works as a spatial reference against which s is compared.

One previous fMRI study suggested the inferior temporal pole region as the locus of c (White et al., 2012). Unfortunately, we could not confirm this finding, because this region was not fully imaged for all participants (see Methods). Nonetheless, their failure of finding neural correlates of c in the IPL or STG could be due to the fact that the effect of past stimuli was not considered in defining c.

### Brain signals of inferred stimulus and decision variable

The s signal in the DLPFC is compatible with its suggested roles in maintaining current sensory information during decision-making (Curtis and Lee, 2010; Haller et al., 2018). The s signal in the cerebellum is also in line with previous studies indicating the involvement of the cerebellum in sensory perception (Baumann et al., 2015; Gao et al., 1996). By contrast, we failed to find any signal of s in the early retinotopic visual areas. Is this failure due to the relatively poor spatial resolution of the fMRI imaging protocol used in the current study? To check this possibility, we revisited one of our previous imaging studies (Choe et al., 2014), where high-resolution fMRI responses were acquired in V1 while observers performed the same task as in the current study, and examined the V1 population activity with a fine retinotopy-based population decoding analysis. However, unlike the DLPFC and cerebellum, where the s signals were regressed onto both external stimuli (Z) and decisions (d) (Fig. 14*A,B*), the V1 activity was regressed only onto external stimuli (Z) but not onto decisions (d) (Fig. 14*C*). This implies, as concluded previously (Choe et al., 2014; Jasper et al., 2019; Lee et al., 2007; Nienborg and Cumming, 2009), that the V1 population activity represents physical stimuli (Z) robustly but its trial-to-trial variability does not seem to be strongly involved in determining the trial-to-trial variability in choice (d).

**Figure 14.**
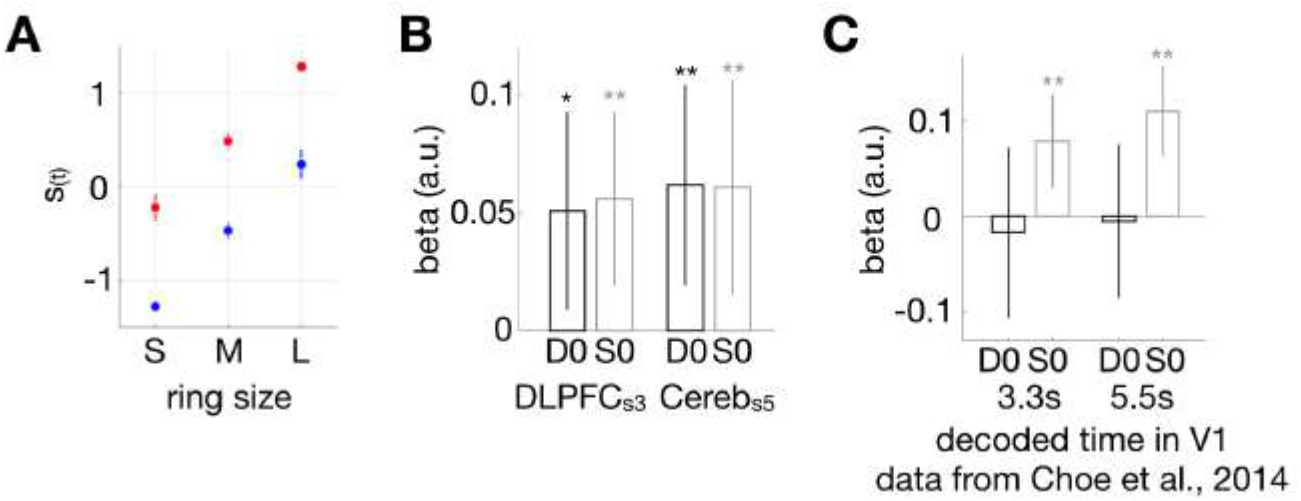
Distinction between brain signal of s and physical ring size. ***A,*** Presence of decision information in the latent variable s of the NBMC. The mean values of s are plotted against the physical ring size, separately for ‘small (blue)’ an ‘large (red)’ decisions. The error bars are 95% bootstrap CIs. ***B,*** GLMM of the brain signal of s in the DLPFC and cerebellum onto current decisions (D0) and current stimuli (S0). ***C,*** GLMM of the brain signal of s in the primary visual cortex onto D0 and S0. The error bars are 95% GLMM CIs, and asterisks indicate p values: *< 0.05; **< 0.01.

## Footnotes

1 We are aware that ‘classification’ and ‘categorization’ are used interchangeably. But we insist that these two tasks, despite their superficial similarity, must be distinguished because they differ in computational architecture. The implications of this distinction will be addressed in Results and Discussion.

## Notes

There is no financial interests or conflicts of interest

#### Summary of Updates

Introduction and Discussion have been re-written based on the comments on the earlier version of the manuscript. The supplementary materials have now been brought to the main text, with additional figures for Results.

